# Evolutionary History of Calcium-Sensing Receptors Unveils Hyper/Hypocalcemia-Causing Mutations

**DOI:** 10.1101/2023.06.11.544489

**Authors:** Aylin Bircan, Nurdan Kuru, Onur Dereli, Berkay Selçuk, Ogün Adebali

**Affiliations:** Faculty of Engineering and Natural Sciences, Sabanci University, İstanbul, Türkiye; TÜBİTAK Research Institute for Fundamental Sciences, Gebze, Türkiye; Department of Microbiology, The Ohio State University, Columbus, Ohio, USA

## Abstract

Calcium-sensing receptor evolution highlights hyper/hypocalcemia-causing mutations The Calcium Sensing Receptor (CaSR) is a key player in regulating calcium levels and has been linked to disorders like hypercalcemia and hypocalcemia. Despite advancements in understanding CaSR’s structure and functions, there are still gaps in our understanding of its specific residues and their differences from receptors within the same class. In this study, we used phylogeny-based techniques to identify functionally equivalent orthologs of CaSR, predict residue significance, and compute specificity-determining position (SDP) scores to understand its evolutionary basis. The analysis revealed exceptional conservation of the CaSR subfamily, with high SDP scores being critical in receptor activation and pathogenicity. To further enhance the findings, gradient-boosting trees were applied to differentiate between gain- and loss-of-function mutations responsible for hypocalcemia and hypercalcemia. Lastly, we investigated the importance of these mutations in the context of receptor activation dynamics. In summary, through comprehensive exploration of the evolutionary history of the CaSR subfamily, coupled with innovative phylogenetic methodologies, we identified activating and inactivating residues, providing valuable insights into the regulation of calcium homeostasis and its connections to associated disorders.

## Introduction

Calcium sensing receptor (CaSR) is a class C G-protein-coupled receptor (GPCR) that maintains extracellular Ca^2+^ homeostasis by sensing calcium ions in the blood and regulating parathyroid hormone release and urinary calcium (Cook *et al*, 2015; Gorvin, 2019). The CaSR is activated by Ca^2+^ and L-amino acids such as L-Phe and L-Trp as well as polyamines and polypeptides (Chen *et al*, 2021; Geng *et al*, 2016; Ling *et al*, 2021). Like the other class C GPCRs such as metabotropic glutamate receptors, ligands bind to the extracellular Venus flytrap (VFT) domain of the receptor (Wootten *et al*, 2018).

Class C GPCRs are obligate dimers, forming either homo or heterodimers (Wootten *et al*., 2018). CaSR forms a homodimer where each subunit is composed of an extracellular domain (ECD), comprising a bilobed (LB1, LB2) VFT and a cysteine-rich domain (CRD) connected to a heptahelical transmembrane (7TM) domain (Chen *et al*., 2021; Ling *et al*., 2021).

Crystal structures of the ECD (Geng *et al*., 2016; Zhang *et al*, 2016) and cryo-electron microscopy structures of the full-length CaSR (Chen *et al*., 2021; Gao *et al*, 2021; Ling *et al*., 2021; Park *et al*, 2021a; Wen *et al*, 2021) reveal the structural basis for activation mechanisms and ligand binding sites. L-amino acid binding site at the interdomain cleft of LB1-LB2 (Chen *et al*., 2021; Geng *et al*., 2016; Ling *et al*., 2021; Mun *et al*, 2005; Zhang *et al*, 2014; Zhang *et al*, 2002) and multiple Ca^2+^ amino acid binding sites on the VFT domain are shown in the literature (Chen *et al*., 2021; Geng *et al*., 2016; Ling *et al*., 2021). While Ca^2+^ is the composite agonist for the CaSR, L-amino acids promote receptor activation along with Ca^2+^, but they are not able to activate the receptor alone (Liu *et al*, 2020). Even though Ca^2+^ alone activates the receptor in functional assays (Liu *et al*., 2020), whether it activates the CaSR in the absence of L-amino acid is still controversial (Chen *et al*., 2021; Ling *et al*., 2021).

Variants in CaSR may cause malfunctions that result in Ca^2+^ homeostasis diseases in humans. More than 400 germline loss/gain-of-function (i.e., LoF and GoF, respectively) mutations cause hypercalcaemic disorders, neonatal severe hyperparathyroidism (NSHPT), familial hypocalciuric hypercalcemia type-1 (FHH1), and autosomal dominant hypocalcemia type-1 (ADH1), respectively (Gorvin, 2019). Many more CaSR variants are anticipated to be identified as more population-level genetic data become available (Gorvin, 2019). Gaining insight into the function of individual residues within the receptor structure and their involvement in activation mechanisms has the potential to enhance our understanding of the probability of variant pathogenicity and the signaling processes of the CaSR. The examination of receptors within a family and across different families allows for the identification of the specific function of each residue in a receptor. However, the comprehensive understanding of the structure and activation mechanisms of several families within the class C GPCRs remains elusive. This is particularly true for the G-protein coupled receptor family C group 6 member A (GPRC6A) and the type 1 taste receptors (TAS1Rs; specifically members 1, 2, and 3), which are the most closely related subfamilies to the CaSR.

While all subfamily receptors of class C GPCRs share common domains and structural features, the details of responding to different ligands and activating signaling pathways may differ even between closely related receptors (Wootten *et al*., 2018). Gene duplication is the main mechanism that generates new protein functions across GPCRs (Flock *et al*, 2017). Protein families are evolved by speciation events following gene duplication (Chagoyen *et al*, 2016; Studer *et al*, 2013); thus, sequence comparisons of members within a subfamily and between subfamilies can show the evolutionarily conserved domains as well as diverged protein sites that distinguish one subfamily from others. One challenge with this analysis is that excessive gene duplication events complicate the identification of functionally identical orthologs in a subfamily. Moreover, the conservation patterns in paralogs and distant homologs may help infer the specific roles of a single residue in protein function. Because the evolutionary pressure on paralogs and close orthologues is not the same, substitutions allowed in paralogs may not be acceptable in close orthologues. Thus, using functionally identical orthologs in sequence comparisons is crucial to inferring the role of each residue in a protein family.

Here, we show the importance of each residue in CaSR by comparing it with the closely related subfamilies, GPRC6A and TAS1Rs. We identified all orthologous sequences in each subfamily by building phylogenetic trees and manually curating the duplications/speciations on the tree to obtain all functionally equivalent orthologs within each subfamily. To obtain members of a subfamily without requiring computationally expensive phylogenetic tree building and manual curation steps, we generated highly sensitive subfamily-specific profile hidden Markov models (HMMs) by using the functionally equivalent orthologs we determined using phylogenetic tree analysis. We calculated a specificity score for each residue in a subfamily by calculating scores based on a modified version of the PHACT algorithm (Kuru *et al*, 2022) scores which considers independent evolutionary events on the phylogenetic tree while scoring the acceptability of an amino acid substitution. We predicted the functional consequence of every potential substitution in CaSR by using the gradient boosting trees machine learning approach. Lastly, we investigated how our predictions relate to the activation mechanism of CaSR.

## Results

### Evolutionary History of Class C GPCRs

To reveal the evolutionary constraints on protein families, we developed a strategy to precisely define a protein subfamily. A precise subfamily definition can be achieved by revealing the evolutionary history of the superfamily. The evolutionary history of gene families can only be established by reconstructing high-quality phylogenetic trees, which can be used to pinpoint gene duplication events. Discrimination between gene duplication and speciation nodes enabled us to define the paralogous and orthologous protein sequences. We further analyzed the phylogenetic trees to classify the orthologous sequences that are likely equivalent in function. A subfamily is defined by the human receptor and its orthologs. Within class C CPCRs, there are twenty-two distinct subfamilies: CaSR, GPRC6A, three taste receptors (TAS1R1-3), eight metabotropic glutamate receptors (mGluR1-8), GPRC5A-D, GABBR1, GABBR2, GPR156, GPR158, and GPR179. We used functionally-equivalent orthologs in comparative analyses between subfamilies, which eventually yielded subfamily-specific signatures that can be used to define that particular subfamily and its function. Finally, the association between the signature and function would enable a better understanding of specific molecular mechanisms and the effects of variants, particularly for the protein subfamily of interest. Here, we aim to reveal the signatures of the CaSR subfamily that is implied in the specific function of calcium-sensing and downstream signaling.

We have retrieved the complete proteomes of 478 species from the NCBI database. To identify proteins that belong to the class C GPCR family, we performed a hmmsearch using the seven transmembrane domain profile (Pfam: 7tm_3) (Fig 1) against the proteomes. While this search allowed us to retrieve the entire class C GPCRs hitting the hmm profile, it did not provide subfamily annotations for the 22 subfamilies in class C GPCRs. To select canonical isoforms, we performed a profile hmmscan of the PfamA profile against Class C GPCR. To generate a general HMM profile for each subfamily, we first applied a BLAST search using each human class C GPCR as a query (Altschul *et al*, 1990). For each subject, we blasted them against the human proteome and retrieved the bidirectional best hits (Core subfamily assignment, Fig. 1).

**Figure 1:**
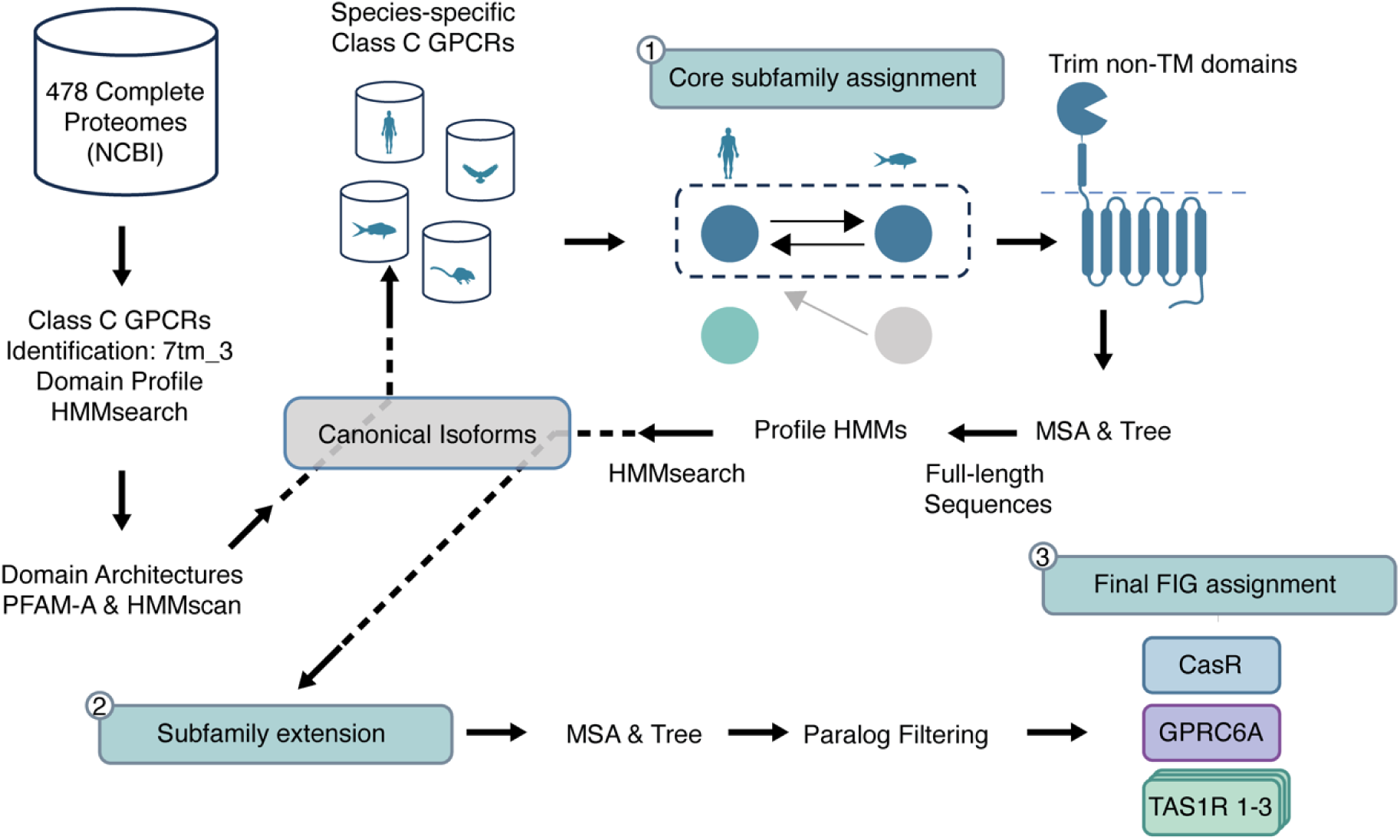
Summary of the Methodological Framework. 478 complete proteomes were retrieved from the NCBI database. Each sequence was searched by hmmsearch against the Pfam 7tm_3 domain profile to retrieve all class C GPCRs. Domain architectures of class C GPCRs were determined by hmmscan against Pfam. A profile to identify canonical isoforms. Species-specific BLAST databases of the canonical isoforms were built. Bi-directional mutual best hits were detected by blasting each canonical sequence against the species databases (core subfamily assignment). TM domains of core subfamily sequences were aligned, and ML trees were built to make subfamily profile HMMs. By hmmsearch against subfamily profile HMMs, other sequences in the subfamilies were found (subfamily extension). Sequences in each subfamily were aligned, and ML trees were built. Based on the ML trees, paralogs were filtered, and functionally identical groups were identified (FIG).

For proteins that did not have bidirectional mutual best hits, we assigned them to a subfamily based on their homology search against the HMM profiles generated in the previous step (Subfamily Extension, Fig 1). We produced maximum likelihood (ML) trees of extended subfamilies and filtered paralogous sequences to obtain functionally identical groups (FIGs).

The CaSR subfamily produced over five thousand hits, which included vomeronasal and olfactory receptors that have never been shown to sense calcium. Previous research has shown that CaSR is classified in the pheromone/olfactory cluster of class C GPCRs (Pin *et al*, 2003). In species that had multiple proteins assigned to the CaSR subfamily, we constructed a ML tree using these hits and other human class C GPCR protein sequences. These trees revealed that a significant number of duplication events occurred in the species after the clade diverged from CaSR. As a result, we defined this diverged clade as a new subfamily named CaSR-likes. The sequences in this subfamily are unlikely to maintain calcium homeostasis, and therefore should not be annotated as calcium-sensing receptors.

We selected representative sequences from different species for each subfamily of 22 different receptor subfamilies and 264 CaSR-like sequences and built a ML tree (Fig 2A). Also, we built the ML trees of all proteins from CaSR, GPRC6A, taste receptors and merged these trees to the representative tree of class C GPCRs (Fig 2A). The resulting phylogeny shows that are five major clades: CaSR-related, GABA, mGluR, orphans, and retinoic acid-induced (RAIG). Orphan receptors, GPR158 and GPR179, formed a clade that was diverged from other receptors consistent with previous trees (Harpsoe *et al*, 2017) and had a 0.95 transfer bootstrap expectation (TBE) value. γ-aminobutyric acid-B receptors (GABBR1 and GABBR2) formed another clade diverging from GPR156 with 0.97 TBE. γ-aminobutyric acid-B receptorsreceptors evolved earliest and have a common ancestor with the highest taxonomic rank (33213−Bilateria) compared to other subfamilies. The CaSR group (CaSR, CaSR-likes, GPRC6A and taste receptors) was diverged from metabotropic glutamate receptors (mGluR1-8) and RAIG receptors (GPRC5A, GPRC5B, GPRC5C, GPRC5D) with 1 and 0.98 TBE values, respectively. Within the CaSR group, clade CaSRs and CaSR-likes were diverged from GPRC6A and taste receptors with 1 TBE. Except for TAS1R1 and TAS1R2, all CaSR group subfamilies have a common ancestor from taxonomy clade 7776-Gnathostomata. TAS1R1 and TAS1R2 were more specific than other CaSR group subfamilies that evolved from 117571-Euteleostomi. Comparison analysis of branch lengths (Patil, 2021) among common species between CaSR, GPRC6A and taste receptors shows that the CaSR subfamily is significantly more conserved than its closest subfamilies (Fig 2B)

**Figure 2:**
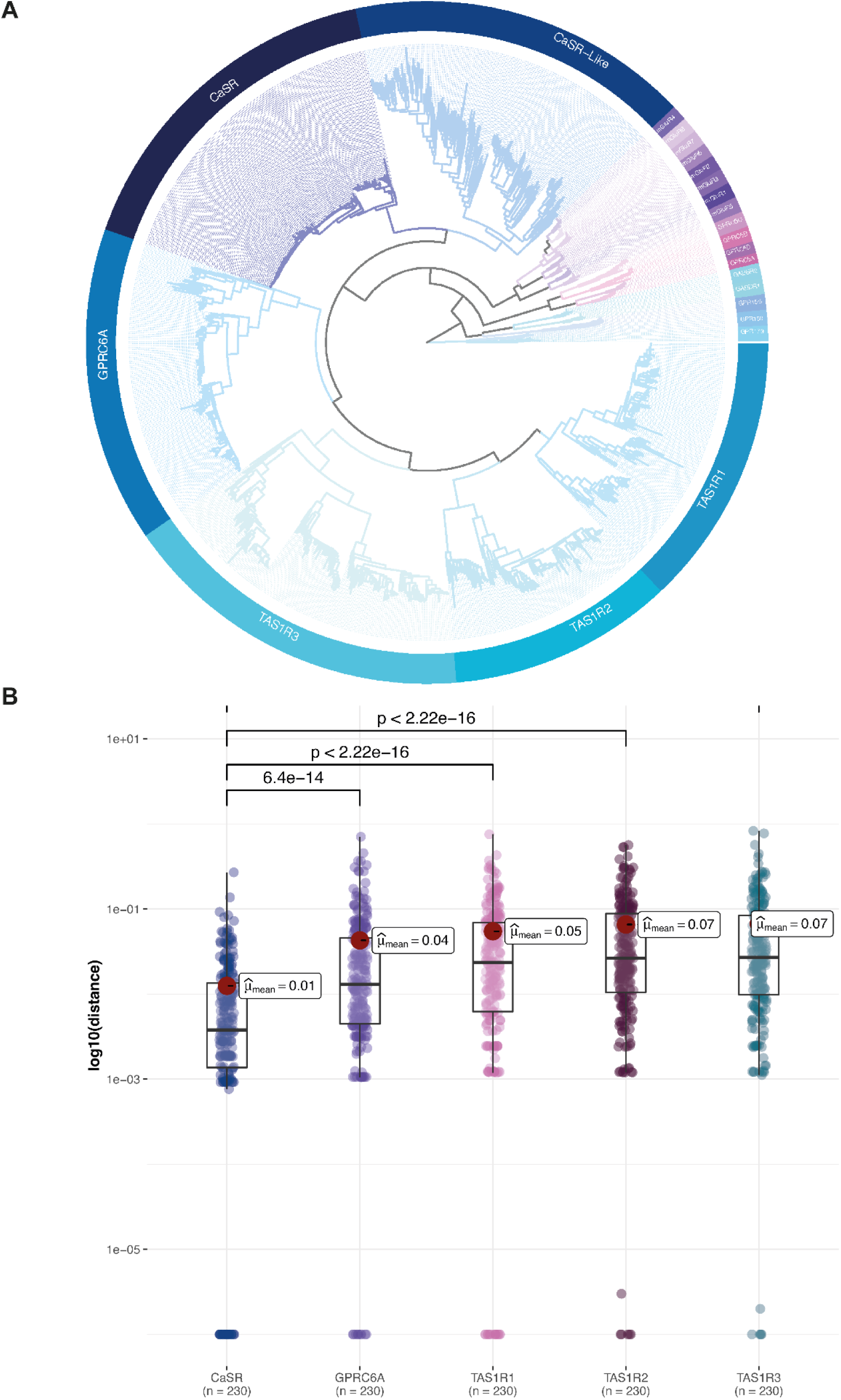
Evolution of Class C GPCRs. **(A)** The maximum likelihood phylogenetic tree of Class C GPCRs, spanning representative species from each subfamily, is shown. Subfamilies are represented as circular layers around the ML tree. All twenty-two Class C GPCR subfamilies are shown in the inner circle. In addition to these subfamilies, vomeronasal and other orphan receptors are represented as CaSR-like receptors. All proteins in CaSR, GPRC6A and TAS1Rs are merged into this representative species tree. **(B)** Branch lengths from leaf to root of the common species that exist in all CaSR, GPRC6A and TAS1Rs are taken from the subfamily trees. Welch’s t-Test using the ggstatsplot package results are shown on the graph.

The higher diversity of CaSR-likes relative to CaSRs is reflected in the ML tree (Fig 2A). Branch lengths of CaSR-likes are longer in contrast to shorter branch lengths in CaSR. Longer branch lengths show that more variation, and thus divergence*, occurred in the CaSR-like clade. Moreover, extensive gene duplication events occurred in this clade. For instance, rodents such as Dipodomys ordii (taxid: 10020),* Octodon degus (taxid:10160) and snakes such as Notechis scutatus (taxid: 8663) have more than a hundred receptors matching the CaSR profile. However, these matches include type 2 vomeronasal receptors (V2R) and V2R-likes. Among mammals, V2R genes exhibit significant variation. While dogs, cows, and primates except prosimians do not have functional V2Rs, rodents, reptiles, and fish have multiple intact V2Rs (Goes van Naters & Mucignat-Caretta, 2014). Since these receptors do not have functional orthologs in mammals, they are likely to be functionally diverged, and it is crucial to separate them from functionally-equivalent CaSRs.

#### Subfamily-specific Profile HMMs to Obtain Orthologs

In the class C GPCR family, gene duplication events give rise to new specificity, and each duplicated gene with a new function evolves through further speciation events, producing a set of orthologous sequences (Chagoyen *et al*., 2016; Studer *et al*., 2013). Each subfamily of class C GPCRs shares a relatively conserved membrane-spanning region but also exhibits a degree of variability that underlies functional differences. At the molecular level, residues that are responsible for certain functional characteristics such as ligand and coupling selectivity are called specificity-determining residues (Chagoyen *et al*., 2016). Conservation analysis from multiple sequence alignments (MSAs) can be used to find residues that are conserved in all subfamilies through evolution as well as specificity-determining residues that are only conserved in a subfamily and differ in other subfamilies. However, the success of this method depends on the sequences that are used to build alignments. Therefore, it is vital to use functionally identical orthologs in the analysis.

The seven-transmembrane domains of class C GPCRs are used to build a class-specific general profile for this family (Pfam:7tm_3). However, this domain does not contain enough information to differentiate subfamilies further. Moreover, excessive gene duplication events, as seen in the CaSR-like clade, require precise phylogenetic analysis to differentiate between CaSR and CaSR-like sequences. Also, subfamily specific profile HMMs are shown to be promising methods to detect protein sequences belonging to a protein subfamily, as well as separation of homologs and non-homologs *(Brown et al, 2005; Srivastava et al, 2007)*. In this paper, we present a novel approach to constructing subfamily-specific profile HMMs based on the precisely produced subfamily alignments and trees. The general idea of the approach is given in Figure 3.

**Figure 3:**
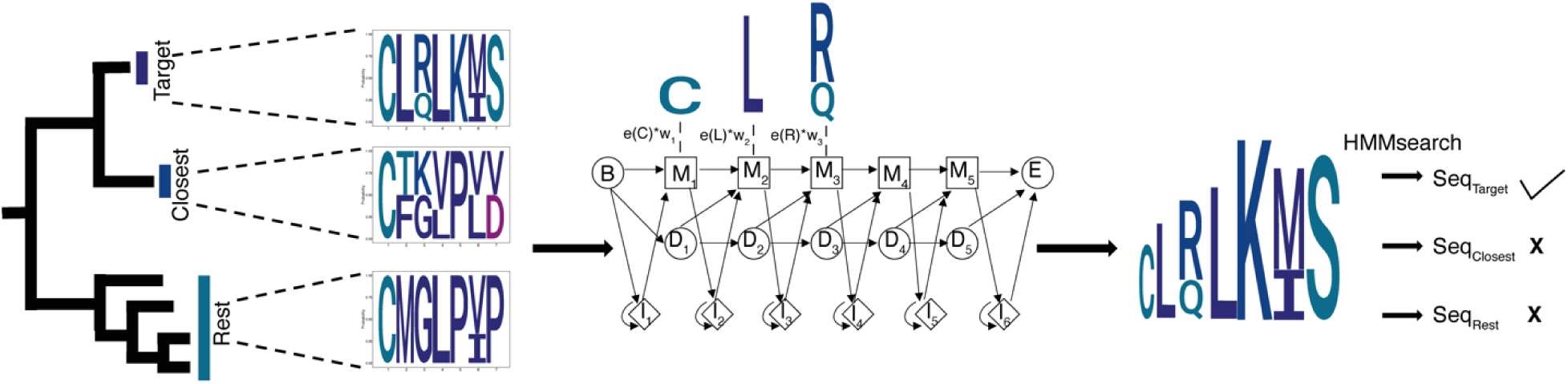
Subfamily Specific HMM Models. Based on the phylogenetic tree, the target, the closest, and the rest groups were determined. Initial HMMs were built without using priors. Representative amino acids in each group are selected, and their scores are calculated. According to groups, representative amino acids, and conservation scores, we calculated weights to change the emission probabilities of initial HMMs.

To construct subfamily specific profile HMMs, we first define the target family, its closest family (phylogenetic neighboring clade), and the rest based on the phylogenetic tree. We weight the identity score of each amino acid to calculate the emission probabilities. The highest weight is given to the residues which are only conserved in the target subfamily; hence, they differentiate one subfamily from the others. A minimum weight is given to the residues that are conserved both in the target subfamily and its closest clade. We tested our subfamily-specific profile HMMs’ performance on independent sequences retrieved from UNIPROT (Roberts *et al*, 2019) which were not used in calculating the position weights. We assigned sequences to their corresponding subfamilies by following the same steps as the NCBI dataset (Pruitt *et al*, 2007) used to build these models. We selected new taxa that were not in the NBCI dataset to test the performance of our profiles, and our subfamily specific profile HMMs correctly identified all members of a subfamily while avoiding hits to proteins from other subfamilies (Table 1).

**Table 1:** Subfamily Specific Profile HMM’s Performance.

| Subfamily HMM | Test Cases | True Positives | False Positive | False Negative | True Negative |
| --- | --- | --- | --- | --- | --- |
| CaSR | 81 | 81 | - | - | 232 |
| GPRC6A | 62 | 62 | - | - | 251 |
| TAS1R1 | 75 | 75 | - | - | 238 |
| TAS1R2 | 21 | 21 | - | - | 292 |
| TAS1R3 | 74 | 74 | - | - | 239 |

### Specificity Determining Residues

CaSR is distinguished from other subfamilies of class C GPCRs by its oversensitivity to substitutions that can result in either GoF or LoF mutations. This sensitivity is due to its critical role in maintaining systemic calcium homeostasis and its high responsiveness to very slight changes in extracellular Ca^2+^ concentrations (Gorvin, 2021). As CaSR is the most conserved and ancestral subfamily among CaSR-likes, GPRC6A and TAS1Rs, it is reasonable to expect that some positions can be under the relaxation of existing purifying selection in CaSR-likes, GPCR6A or TAS1Rs, but not in CaSR itself. Conversely, at some positions, the same amino acid may retain functional significance in both subfamilies, and at others, a position remains important in each subfamily, but different amino acids are favored in each duplicate.

Specificity-determining residues that are conserved in subfamily, but differ from its sister clade can be predicted by directly comparing ancestral family sequences and calculating their divergence scores (Bradley & Beltrao, 2019). However, using MSAs alone does not account for the number of substitution events. For example, a single substitution event in the common ancestor of the bony fish clade of the CaSR subfamily can be inherited into multiple descendants’ sequences. Assessing this single substitution event as it repeats in each sequence independently results in overcounting of these changes. Due to this mistake and overcounting the effect of one single mutation repeatedly, the position is considered (i) to tolerate that particular amino acid and (ii) functionally less important. In contrast, a single evolutionary event might have been compensated by other substitutions in the same evolutionary node. Such a substitution might not be tolerated in the other clades of the subfamily.

Another consideration to identify and order specificity-determining residues is treating substitution events on the phylogenetic tree unequally. When an amino acid in CaSR remains the same but can differ in the nearby CaSR-likes subfamily, it indicates that the amino acid has a CaSR-specific role. We expect the SDP score of such an amino acid to be high compared to others. If an amino acid is conserved in both CaSR and remote subfamilies like taste receptors but likely to be substituted in CaSR-likes, it suggests that the amino acid plays a common functional role in both CaSR and other subfamilies. For such an amino acid, the SDP score must be low, since it is not a specific position for CaSR.

*To consider these two important factors, we designed an approach to identify and prioritize* residues by specificity, which differentiate a subfamily from others in the CaSR group (CaSR, GPRC6A and TAS1Rs). Our approach is based on the idea presented in the functionally divergent residues method (Bradley & Beltrao, 2019) along with adaption of the PHACT method (Kuru *et al*., 2022). We calculated the probability of each amino acid at each node of the CaSR-group phylogenetic tree by ancestral sequence reconstruction (Fig 4A). Starting from the root of the tree, we identified each substitution event and at which subfamily node that event happened. After counting the number of independent substitution events in a subfamily clade and comparing the probability of the same substitution in other subfamily clades, we ordered the specificity-determining residues. We assumed that if an amino acid is allowed to change on sister subfamily nodes and is poorly conserved in sister subfamily nodes while it is highly conserved on the target subfamily node, it is a specific residue to the target subfamily only. If a substitution event is observed on a clade close to the target node, we consider that event to increase the specificity of a residue because it diverges the target group from closest sister clade. The details and the algorithm are given in materials and methods (Algorithm 3).

**Figure 4:**
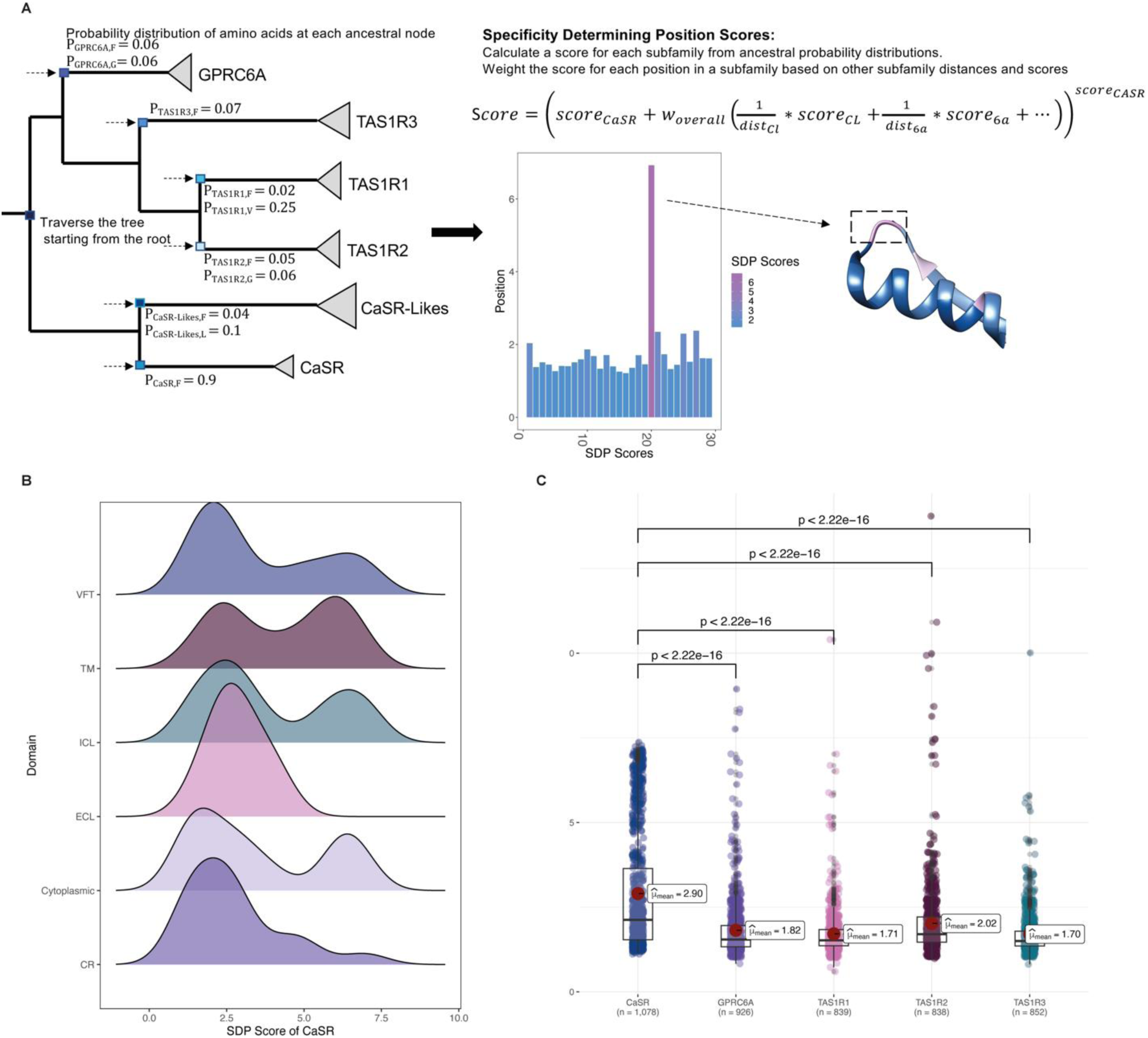
Specificity Determining Position Scores. **(A)** The calculation of SDP scores uses the phylogenetic tree and the probability distribution of amino acids at each ancestral node. **(B)** The SDP score distributions of CaSR among different domains and the SDP score distributions of each subfamily are shown. **(C)** Welch’s t-Test shows that CaSR has more residues with higher SDP scores compared to GPRC6A and TAS1Rs.

We calculated specificity scores for each CaSR, GPRC6A and TAS1Rs. Specificity score distributions show that CaSR has more specific residues compared to other subfamilies (Fig 4B). On the VFT domain, specific residues are clustered in different regions (Fig 5). We found a cluster of specific residues on the interdomain cleft between LB1-LB2 which is the L-amino acid binding site in other class C GPCRs (Chen *et al*., 2021). It suggests that this region is the primary Ca^2+^ binding site in CaSR, consistent with (Liu *et al*., 2020). We found two different clusters of specific residues on the ECD. First cluster was on the LB1 domain and on the LB1-LB1 dimer interface. LB1 domain plays a role in anchoring ligands and initiating domain twisting by conformational changes at the interface between LB1 regions (Chen *et al*., 2021; Ling *et al*., 2021). The second cluster was found at the cytosolic side of the LB2 and at the interface between LB2-CRD, where Ca^2+^ ions bind (Chen *et al*., 2021; Geng *et al*., 2016; Ling *et al*., 2021). Interaction between LB2 subunits is required for CaSR activation that propagates to large-scale transitions of the 7TMDs (Chen *et al*., 2021; Ling *et al*., 2021). Specific residues on the LB1 domain, LB1-LB1 dimer interface, and LB2-CRD interface indicate that they provide the structural conformational changes upon ligand binding to the interdomain cleft. Mutations located in these regions are associated with LoF and GoF mutations (Fig 7B) (Gorvin, 2019). Other specific residues are found on the CR, ECL2 and TM domains. On the ECL2 acidic residues D758 and E759 are specific to CaSR. The intersubunit electrostatic repulsion between the ECL2 regions could facilitate the activation of CaSR (Chen *et al*., 2021; Ling *et al*., 2021). In the agonist+PAM bound state the ECL2 is moved by the interaction among E759, W590, and K601. Deletion of D758 and E759, and single mutations of K601E and W590E disrupt CaSR activity; however, Δ758–759 mutant was expressed at the cell surface with comparable levels to that of WT while W590E and K601E mutants were expressed on the cell surface lower than the WT level (Chen *et al*., 2021). We found that residues W590 and K601 are not specific to CaSR. The TM domains of two protomers of CaSR come into close proximity upon receptor activation (Ling *et al*., 2021).

**Figure 5:**
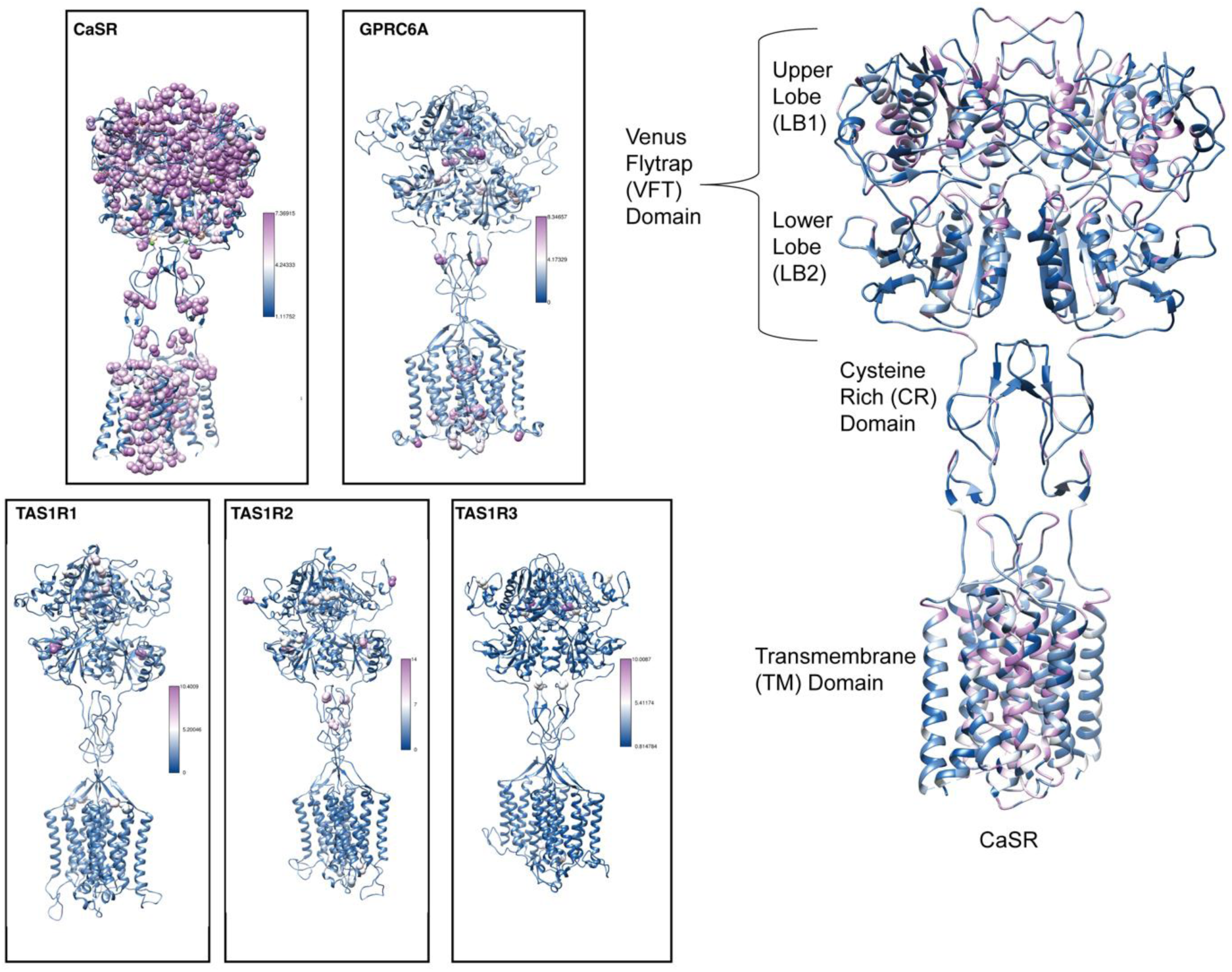
Specific conserved residues mapped onto structures. The cryo-EM structure of human CaSR bound with Ca^2+^and L-Trp (PDB:7DTV) and homology models of GPRC6A, TAS1R1, TAS1R2 and TAS1R3 are colored based on SDP scores. Residues with a high SDP score (above 5.0) are shown as spheres. Domains on the human CaSR structure (PDB: 7DTV) are labeled and colored according to their SDP scores on the right-hand side.

The interaction between TM4-5 of each subunit in the inactive state is essential (Liu *et al*., 2020), while the interaction between TM6-TM6 is crucial for the active state (Chen *et al*., 2021; Gao *et al*., 2021; Liu *et al*., 2020). The structural findings and the presence of CaSR specific residues on each TM domain suggest that CaSR is specialized in both dimerization and ligand binding. Specific residues on the TM domain are likely play a role in the regulation of conformational changes observed during activation upon ligand binding and inactive states. Residues inside the dimerization interface and interacting with the ligand are quite sensitive substitutions because they can induce malfunctions in the receptor easily. On the other hand, GPRC6A and taste receptors are more tolerant to substitutions, and they are not very specialized respond to a single ion. GPRC6A and taste receptors are activated by a broad spectrum of ligands (Chun *et al*, 2012; Pi *et al*, 2017). Even though the ligand of GPRC6A is controversial in the literature, multiple ligands such as osteocalcin (Ocn), testosterone, basic amino acids and cations such as L-Arg, L-Lys, L-Orn, calcium, magnesium, and zinc are suggested to bind GPRC6A (Pi *et al*., 2017). Taste receptors bind to different ligands, including sugar, L- and D-amino acids, sweet proteins, and artificial sweeteners (Nango *et al*, 2016).

On the TM region, we also find the CaSR specific cholesterol recognition/interaction amino acid consensus (CRAC) motif (L783, F789, S820) that is defined by the consensus (L/V)X1–5YX1– 5(R/K) and is often present at junctions between membrane- and cytosol-exposed domains and shown in the mGluR2 receptor (Kumari *et al*, 2013). Phylogenetic analysis shows that TAS1R3 evolved earliest (7776 Gnathostomata) among TAS1Rs, TAS1R1and TAS1R2 subfamilies have a common ancestor 117571 Euteleostomi. TAS1R3 forms heterodimers with TAS1R1 and TAS1R2 (Chun *et al*., 2012; Nango *et al*., 2016; Nuemket *et al*, 2017). Interactions between the cytosolic terminus of the extracellular CRD is needed for T1R3 dimerization. TAS1R1 and TAS1R2 recognize a broad spectrum of L-amino acids that bind to the intercleft between LB1-LB2 and induce the positional shift of the CRD regions; however, T1R3 loses the corresponding function (Nuemket *et al*., 2017). Our analysis showed that TAS1R1 has specific residues on the LB1, LB2 and extracellular loop regions. Also, TAS1R2 has specific residues on the LB1, LB2 and CR domains. On the other hand, in TAS1R3, we found specific residues only on the LB1 and one on the CR domain. Since LB1-LB2 domains create a cavity for ligand binding, specific residues on the LB1-LB2 domains of TAS1R1 and TAS1R2 may contribute to domain transformation upon ligand binding. However, the number and distribution of specific amino acids suggest that taste receptors are not under the same strict selective pressure as CaSR.

### Gradient Boosting Trees Machine Learning Approach to Predict the Mutation Types in CaSR

Because CaSR is a highly conserved subfamily, any substitution on the receptor disrupts the function of the receptor and causes either GoF or LoF mutations. However, predicting the functional consequences of a substitution is challenging. Evolutionary conservation of a residue among subfamilies might reflect the common structural constraints, but it does not distinguish between LoF and GoF mutations. In addition, at certain positions, substitution of different amino acids causes either LoF or GoF mutations (Goes van Naters & Mucignat-Caretta, 2014). We hypothesized that “activating” mutations are more likely to be tolerated in neighboring clades such as GPRC6A and TAS1Rs and not in CaSR, whereas, in general, LoF (inactivating) mutations are not tolerated in the larger clade of these receptor subfamilies. To test this hypothesis and determine we can discriminate between GoF and LoF mutations in CaSR, we present a tree boosting machine learning algorithm, XGBoost (Chen & Guestrin, 2016) that links multiple features such as conservation scores, physico-chemical properties of amino acids, and domain information.

Our algorithm uses sequence-based features, identity scores from MSAs, physico-chemical properties of amino acids, and domain information as input features (Fig 6A). Since we calculated our feature values from the MSAs, we divided our dataset into training, validation and test datasets before we created feature matrices to prevent information leakage. We performed 50 replications with different random splittings of datasets to obtain a more robust model performance.

**Figure 6:**
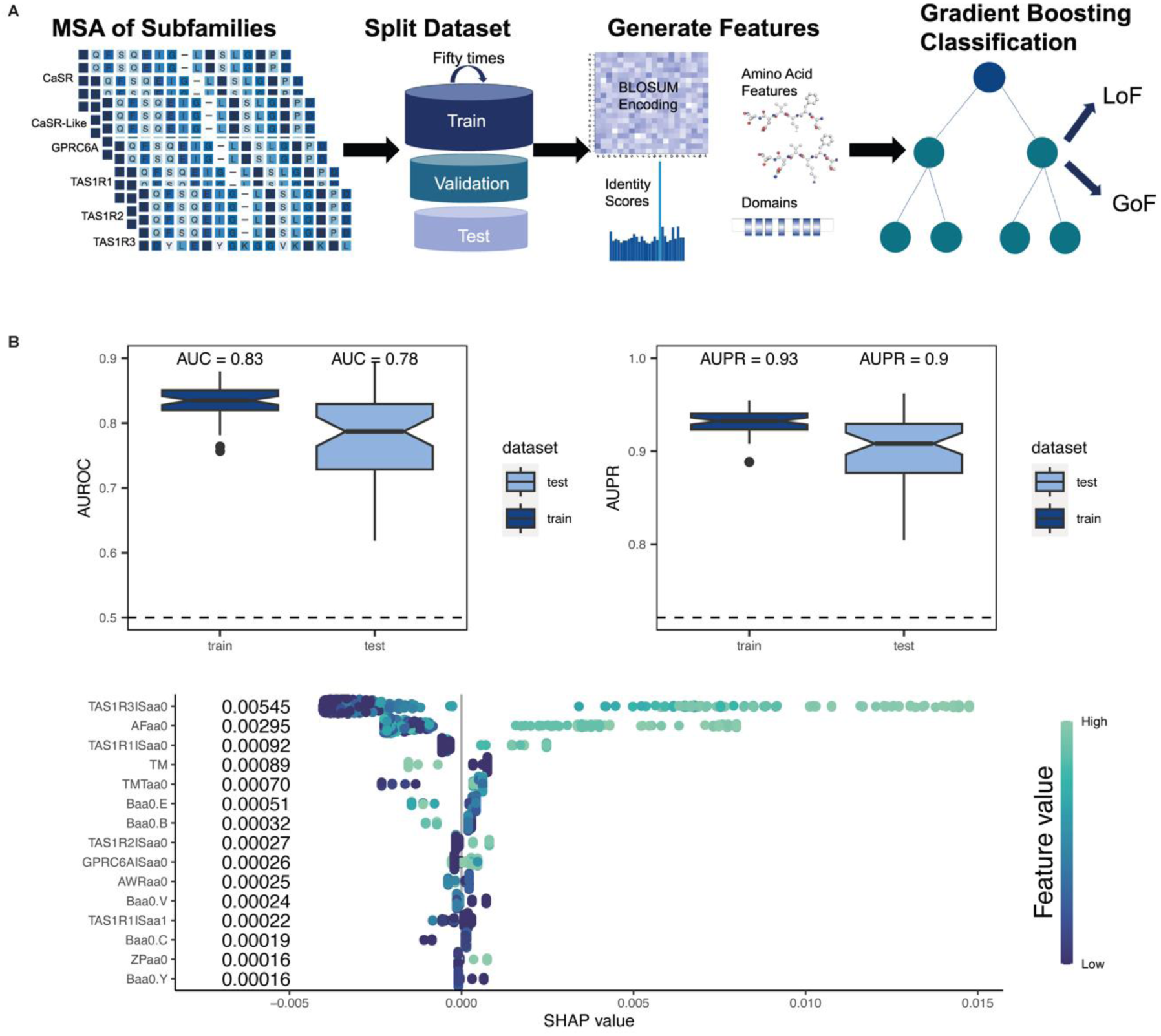
Gradient Boosting Trees Machine Learning Approach to Predict the Mutation Types in CaSR. **(A)** Model architecture. We used MSA of CaSR, CaSR-likes, GPRC6A and TAS1Rs to generate features as well as amino acid physico-chemical features and domain information. We performed 50 replications. **(B)** The performance and feature importance of XGBoost algorithm. The AUROC and AUPR values of 50 replications are shown. The average AUC levels of 50 replications are 0.83 and 0.78 for the train and test respectively. The average AUPR levels of 50 replications are 0.93 and 0.9 for the train and test, respectively. Contributions of Shapley values for type of pathogenicity classification to the model output for XGBoost. aa0: the amino acid found in the human CaSR, aa1: substituted amino acid, AF: average flexibility, TMT: TM tendency, ZP: Zimmerman polarity, B: BLOSUM62, AWR: atomic weight ratio, TM: transmembrane domain

**Figure 7:**
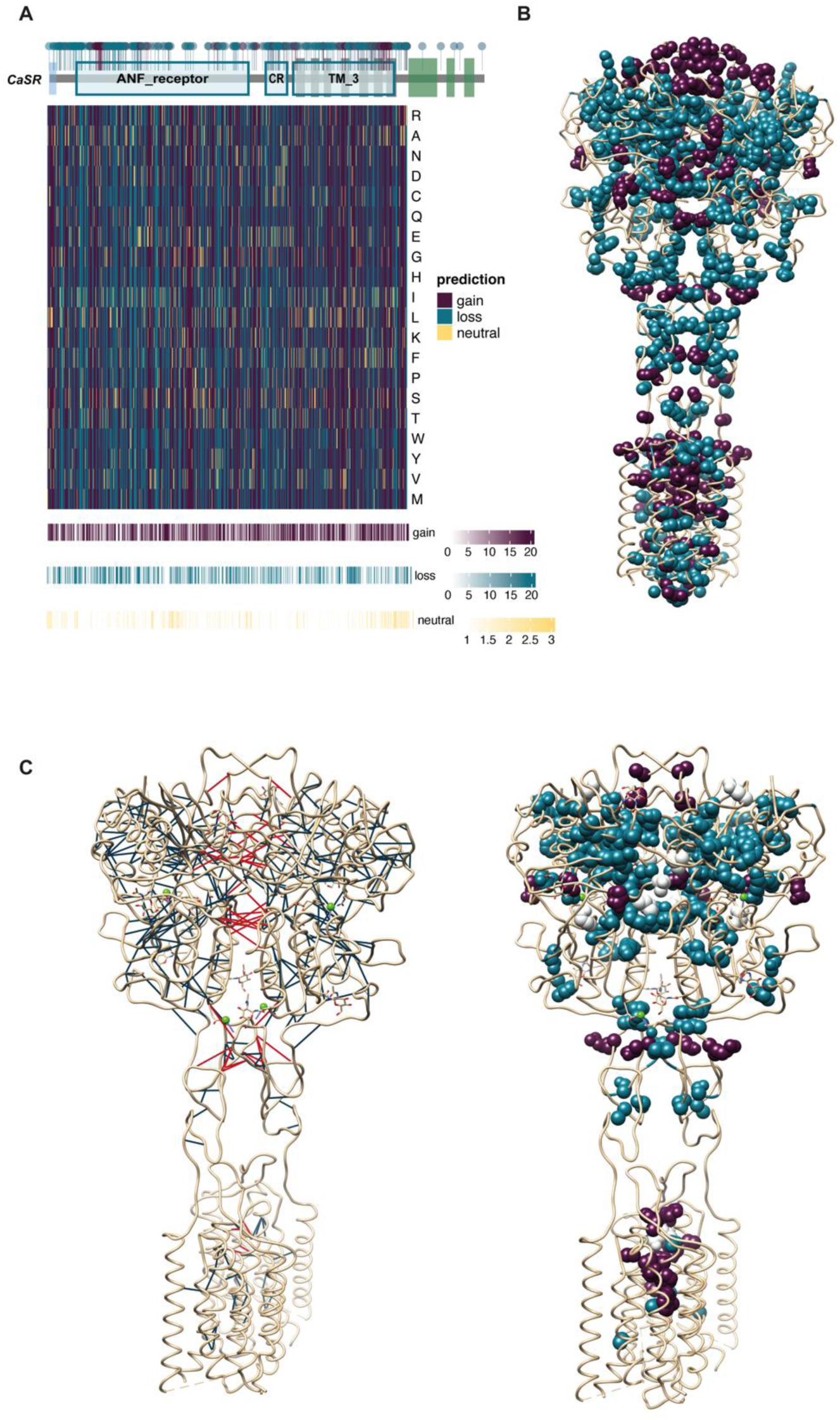
LoF and GoF Mutation Predictions. **(A)** Visualizing the results of our XGBoost model. The heatmap displays the XGBoost model’s predictions for each of the 20 amino acids at every position except disordered regions (892-1078) within the human CaSR. Above the heatmap, the domains of the CaSR are shown. Within these domains, circles represent all known LoF and GoF mutations documented in the literature. Circles denoting GoF mutations are colored purple, while those representing LoF mutations are colored blue. Below the heatmap bar graphs show the number of GoF, LoF and neutral predictions among the 19 possible substitutions. **(B)** Mutations on the human CaSR structure. LoF- and GoF-associated mutations are shown on the cryo-EM structure of human CaSR bound with Ca^2+^ and L-Trp (PDB:7DTV) as blue and red spheres, respectively. **(C)** Increased residue-residue contacts are shown on the cryo-EM structure of human CaSR bound with Ca^2+^ and L-Trp (PDB:7DTV) on the left. Interdomain and intrasubunit interactions are shown as red and blue lines, respectively on the right. LoF- and GoF-associated mutations among the interacted residues are shown as blue and purple spheres, respectively. Switch residues are shown as white spheres.

The ROC and PR curves are used to understand the performance of a binary classifier that assigns each element of data into two groups. The ROC curve is a graphical plot that shows the false positive rate versus the true positive rate for different threshold values between 0 and 1. A PR curve is a plot of the precision and the recall for different threshold values, and it is useful for imbalanced datasets. We used the areas under the ROC and PR curves (i.e., AUC and AUPR, respectively) to compare the performances of the model on the train and test datasets for 50 replications. Higher AUC and AUPR values are associated with better performance. AUC and AUPR over all replications were shown in (Fig 6B). Our average AUC values for training and test among 50 replications are 0.83 and 0.78 (Fig 6B). Our average main AUPR values for training and test among 50 replications are 0.93 and 0.9, respectively (Fig 6B). After we reported our algorithm performance, we trained our algorithm with the whole dataset. We tested our algorithm with new test cases from the literature (Table 2). Additionally, we categorized amino acids that are observed in the CaSR MSA as neutrals. To date, no pathogenic substitution has been reported in the literature for these amino acids that we identified as neutral. We visualized all predictions in the form of a heatmap for every other amino acid at each position until the disordered region (position 892) of the human CaSR (Fig 7A). We mapped known CaSR LoF and GoF mutations on the cryo-EM structure of human CaSR bound with Ca^2+^ and L-Trp (PDB:7DTV)(Ling *et al*., 2021) (Fig 7B). There is a tendency that GoF mutations are in the inner-core regions. In the heatmap, we observed a similar prediction pattern: GoF predictions are mostly in the inner-core regions (Fig 7A). SHAP (SHapley Additive exPlanations) values provide a way to decode the inner workings of a machine learning model like XGBoost. These values calculate the average contribution of each feature to the overall prediction, taking into account any interactions between the features. Based on the SHAP values, the conservation scores of human CaSR amino acids in other subfamilies play a significant role in the model’s prediction, as shown in Figure 7B. If the amino acid is also conserved in GPRC6A and taste receptors (in fact, conservation score in TAS1R3 has the highest contribution), the model predicts a substitution of that amino acid as LoF. On the contrary, when an amino acid is conserved exclusively in CaSR, substituting that amino acid is predicted to result in a GoF. New test cases (Y825F, A840V, L696V, I139T) mentioned in Table 2 have been accurately predicted to cause GoF mutations. These specific amino acids are conserved either solely in CaSR or in GPRC6A and CaSR-like receptors, but not across all receptor types. Notably, the A840V substitution leads to a GoF mutation, as valine is conserved in GPRC6A. These analyses reinforce our hypothesis that residues conserved in CaSR, but not in other subfamilies, are more likely to induce GoF mutations. Additionally, amino acids conserved within closely related subfamilies might disrupt the functioning of other subfamilies. On the other hand, the new LoF cases shown in Table 2 (D99N, C60G, Q164R, T808P) show that reference amino acids are conserved across subfamilies.

**Table 2:** Model’s predictions for the new CASR GoF and LoF mutations from literature. The correct predictions are indicated by a star symbol (*) next to them.

| Mutation | Cause | Prediction |
| --- | --- | --- |
| p.I857S (Chou *et al*, 2022) | hypocalcemia | gain-of-function* |
| p.Y825F (Moon *et al*, 2021) | hypocalcemia | gain-of-function* |
| p.P393R (Palmieri *et al*, 2021) | hypercalcemia | loss-of-function* |
| p.C60G (Li *et al*, 2021) | hypercalcemia | loss-of-function* |
| p.D99N (Hao *et al*, 2021) | hypercalcemia | loss-of-function* |
| p.T186N (Tsuji *et al*, 2021) | hypocalcemia | loss-of-function |
| p.A840V (Roberts *et al.*, 2019) | hypocalcemia | gain-of-function* |
| p.S448P (Dharmaraj *et al*, 2020) | hypercalcemia | loss-of-function* |
| p.L696V (Gomes *et al*, 2020) | hypocalcemia | gain-of-function* |
| p.D433Y (Magno *et al*, 2020) | hypercalcemia | loss-of-function* |
| p.S147L (Majumdar *et al*, 2020) | hypercalcemia | loss-of-function* |
| p.D398N (Zajickova *et al*, 2020) | hypercalcemia | loss-of-function* |
| p.K805R (Sagi *et al*, 2020) | hypercalcemia | gain-of-function |
| p.C60Y (Dong *et al*, 2020) | hypercalcemia | loss-of-function* |
| p.L606P (Wejaphikul *et al*, 2023) | hypercalcemia | loss-of-function* |
| p.H41R (Courtney *et al*, 2022) | hypercalcemia | gain-of-function |
| p.A110D (Bletsis *et al*, 2022) | hypercalcemia | gain-of-function |
| p.I139T (Wu *et al*, 2022) | hypocalcemia | gain-of-function* |
| p.Q164R (Coughlan *et al*, 2022) | hypercalcemia | loss-of-function* |
| p.T699N (Mullin *et al*, 2022) | hypercalcemia | gain-of-function |
| p.R701G (Mullin *et al.*, 2022) | hypercalcemia | loss-of-function* |
| p.T808P (Mullin *et al.*, 2022) | hypercalcemia | loss-of-function* |

Another important feature is the structural location of the amino acid. Our findings indicate that if the amino acid is located in the TM domain, a substitution would result in a GoF mutation. It is known that the majority of GoF mutations are located in the TM domain, as shown in Figure 7B. The presence of certain amino acids on the TM domain of CaSR suggests that they play a crucial role in its activation mechanism. Even though substituting those amino acids might be acceptable in GPRC6A and taste receptors, they might lead to the lock of TM domains and result in the overactivation of CaSR.

To gain insight into the activation mechanism of the CaSR and its association with activating and inactivating mutations, we analyzed the complex network of residue-residue connections by comparing active state CaSR structures with inactive state structures. For the monomer activation network, we modified the residue-residue contact score (RRCS) algorithm (Zhou *et al*, 2019) to detect atom level contact changes in identical amino acids in different states. We made this change for two main reasons. First, class C GPCRs do not exhibit large structural changes during receptor activation (Hauser *et al*, 2021), like other GPCR classes. Therefore, residue level changes do not provide enough resolution for understanding the mechanism, especially for the TM domain. Second, residue level changes can only highlight certain important positions while investigating atom level contact changes enables us to understand the impact of more residues. For the changes observed within the dimerization interface, we used the original algorithm. To build the networks, we identified significant contact changes observed in two receptor states during activation both in the individual protomer and at dimer interface. (Fig 7C). Significantly, the residues that were linked to changes resulting in increased activation were more abundant in critical locations such as the loop of lobe 1 inside the VFT domain, the interdomain cleft, the CR domain, and the TM domains (Fig 7C).

Recent studies have highlighted the role of domain twisting in the CaSR homodimer’s activation process, initiated by intersubunit domain contacts and conformational shifts at the interface between the lobe 1 regions (Ling et al., 2021). In this study, we have noticed a notable augmentation in the level of contact between two pivotal amino acid residues, L125 and Y20, located within the loop region of LB1 domains in the homodimeric structure. Notably, the substitutions of L125F and L125P led to mutants that exhibited a GoF phenotype. In the TM6 domain, we observed increased interaction between residues in the dimer, with particular attention to A824 and P823. Significantly, residue A824, which exhibits specific conservation, has been associated with GoF mutations in the A824P and A824S (Gorvin, 2019)

We identified a notable increase in the level of contacts observed between residues within the CR domain. Specifically, these interactions have been observed between two subunits of the dimer and involve the following interactions: E556-D578, E556-S580, and E556-K552. Although only residue S580 demonstrated specificity for CaSR, other acidic residues either exhibited tolerance towards replacements with other acidic residues or remained conserved across all subfamilies of CaSR. Nevertheless, it is important to acknowledge that the substitution of E556K resulted in a mutation with enhanced activation. The results indicate that the interactions between these particular residues are of utmost importance in maintaining the receptor’s active conformational state. This conclusion is consistent with previous structural investigations that have demonstrated the convergence of LB2, CR, and TM domains in both subunits during the twisting of CaSR, leading to a more condensed conformation in the active state of CaSR (Chen et al., 2021; Ling et al., 2021). In contrast, we have also observed distinct residues linked to LoF mutations. These residues exhibit enhanced interactions within a single subunit, predominantly located in the LB1 and LB2 sections of the VFT domain, specifically in proximity to the Ca^2+^ binding sites (Fig 7C). Residues that induce LoF upon mutation are primarily located within the core of the VFT domain. This implies that any modification in amino acids could potentially induce structural alterations, ultimately resulting in misfolding or disruption of the activation mechanism and consequent LoF.

## Discussion

In this study, we showed the evolution of CaSR by developing a methodology for precisely defining functionally equivalent orthologous sequences across species and therefore subfamilies. We built a high-quality phylogenetic tree of CaSR with its closest subfamilies, GPRC6A and TAS1Rs. Statistical analysis of branch length distances from this phylogenetic tree showed that CaSR is evolutionarily more conserved compared to GPRC6A and TAS1Rs. While GPRC6A and taste receptors can bind to a diverse range of ligands and are able to tolerate substitutions at most of the positions, CaSR requires a delicate balance for proper functioning.

The high evolutionary conservation and specificity of CaSR in contrast to the closest subfamilies are reflected in the SDP score analysis. CaSR has specific residues clustered in different regions of the receptor. They are located on Ca^2+^ and L-Trp binding sites on the VFT, as well as on the dimerization sites between two sub-units of the homodimer. Specific residues on the dimer interfaces indicate that dimerization maintained by interactions between different subunits is required for ligand binding and the correct activation of the CaSR. Ca^2+^ ion binding and interactions between LB2-CR domains and conformational changes in LB1 domain were suggested to be required to activate CaSR (Chen *et al*., 2021; Geng *et al*., 2016; Ling *et al*., 2021). Mutational analysis at some positions on the LB1 domain has been shown to reduce the effect of Ca^2+^-stimulated intracellular Ca^2+^ mobilization in cells (Chen *et al*., 2021; Ling *et al*., 2021). In contrast, substitutions caused negative charge neutralization on the ECL2 result in prompting the activation of CaSR (Ling *et al*., 2021). Our results suggest that residues with low SDP scores on any domain are required for a common activation mechanism since they are conserved across functionally different receptor subfamilies. However, residues with high SDP scores cause malfunctions in the CaSR. Any substitution in a residue with a high SDP score might either cause over or less activation. Deep mutational scanning approaches or new methods that simultaneously profile variant libraries (Jones *et al*, 2020) are needed to provide further evidence to functionally assay all possible missense mutants.

To predict the functional consequence of a mutation in human CaSR, we used the Extreme Gradient Boosting (XGBoost) method. XGBoost is able to perform well on small datasets by incorporating a variety of regularization methods to control the model complexity, which helps to prevent overfitting. We have a small and unbalanced dataset in that the number of GoF mutations was very low, therefore it is prone to overfit. To prevent overfitting while achieving high predictive performance, we used a simple method along with regularization parameters. Moreover, we tried to keep the ratio between the number of LoF and the number of GoF mutations for training and test sets as close as possible. To get a robust performance, we repeated the train-validation-test splitting procedure fifty times. To increase predictive performance, we could use more complex methods, such as deep learning, but they require larger datasets. Studies that used deep learning or ensemble methods for similar assessments are different in terms of prediction, in which they predict the type of mutation as either pathogenic or neutral (Alirezaie *et al*, 2018; Ioannidis *et al*, 2016; Rentzsch *et al*, 2019; Rogers *et al*, 2018). Even though there are a number of mutations of human CaSR in the Clinvar, the functional consequences of most of them are not known.

Given the constraints of the small dataset and limited additional data, we carefully selected and processed the features for our model’s training. Features that are used to train a machine learning model heavily determine its performance. The more features we use, the more information the model has to learn from, which can lead to improved predictive performance. However, having too many features can also lead to overfitting. Moreover, the quality of the features is more important than the quantity. One important evolutionary process that can affect the functional consequences of a substitution is co-evolution. From the MSA of CaSR proteins, we manually selected six positions, p.180,p.212,p.228,p.241,p.557 and p.883, that are in our dataset and co-evolved. We masked the co-evolved amino acids from the MSA and performed the train-validation-test splitting procedure fifty times again. Our average AUC values for training and test among 50 replications were 0.83 and 0.77, respectively and our average AUPR values were 0.93 and 0.89. Despite not experiencing an improvement in performance, we found that the amino acid changes p.I212T, p.F180C, and p.I212S were now predicted to cause LoF, contrary to their previous prediction of causing GoF. We cannot accurately assess the impact of co-evolution on performance because there is a lack of effective tools for identifying co-evolved positions and our dataset contains only a limited number of co-evolved positions, but we anticipate that it is an important feature to differentiate GoF and LoF mutations.

We built subfamily-specific profile HMMs to get all functionally-equivalent orthologs while excluding other proteins. To generate these HMM models, we manually decided target, closest and rest groups based on the phylogenetic tree of CaSR group. Based on the nature of a phylogenetic tree, the selection of these groups is changed, so that this process can be further automated. We did not anticipate that our specific models would match any receptors from other classes of GPCRs, since they are evolutionarily more distant to CaSR group. We expect that our subfamily specific profile HMMs can be used to obtain orthologs in different protein families for the upcoming genomes. They can be particularly useful for studying protein families with many duplications and orphan protein families, where it can be difficult to identify true members. These models are particularly important to avoid computationally expensive and expertise-required phylogenetic tree reconstruction and analysis.

## Materials and methods

### Class C Proteins and Their Domain Architectures

478 complete eukaryotic proteomes were downloaded from the NCBI genomes website (https://ftp.ncbi.nlm.nih.gov/genomes/archive/old ref seq/) in 2018. A hmmsearch of HMMER software (Eddy, 2011) (http://hmmer.org/) was run for each proteome against the Pfam 7tm_3 profile (Finn *et al*, 2016). Sequences with significant 7tm_3 hit based on hmmsearch results (above the default threshold) were compiled from proteomes. A hmmscan of HMMER software (Eddy, 2011) (http://hmmer.org/) was run for these sequences against the Pfam-A 32.0 database (Finn *et al*., 2016). Based on the results of the hmmscan, the longest isoform was taken and saved in a separate file named by taxonomic ID, however, canonical sequences were obtained for human (based on the given canonical proteins on the UniProt website (Consortium, 2017). Because plants do not have GPCRs, they were eliminated from the analysis. For single isoform sequences of each proteome, a BLAST database was built (Altschul *et al*., 1990).

### Subfamily Definition and Subfamily Specific Models

Each protein sequence of each taxon was queried through BLASTP against each prepared BLAST database (Altschul *et al*., 1990). reciprocal mutual best hits of each human class C GPCR were collected in a file named gene id. reciprocal mutual best hits of each class C GPCR and remaining human class C GPCRs were collected and 7TM domains of these sequences were taken based on hmmscan results (the longest sequence that hit the 7tm_3). Sequences were aligned using the MAFFT v7.221 E-INS-I algorithm with default parameters (Yamada *et al*, 2016). A maximum likelihood based phylogenetic tree of each subfamily of class C GPCR was built using RAxML version 8.2.12 with automatic protein substitution model selection (PROTGAMMAAUTO) and 100 rapid bootstrapping parameters (Stamatakis, 2014). The most common lowest taxonomic level was added to the phylogenetic tree with the ETE toolkit (Huerta-Cepas *et al*, 2016). Based on the phylogenetic tree, sequences belonging to the corresponding subfamily were taken, and a profile HMM was built. Subfamily Assignment The process begins by scanning each sequence with a 7tm_3 domain against profile Hidden Markov Models (profile HMMs). After the sequence is scanned, the subfamily is determined based on three conditions: (1) The maximum score value of the hmmscan must belong to the given subfamily. (2) E-value is a measure of the significance of a match in a database search, and the lower the E-value, the more significant the match is. The E-value of the sequence must be the lowest. (3) The sequence must belong to the most common highest taxonomic level of the given subfamily. Taxonomic level refers to the classification of an organism within a biological classification system. If a sequence meets these three conditions, it is assigned to the corresponding subfamily. After this, the full length sequences of each subfamily were then aligned using the MAFFT v7.221 algorithm (Yamada *et al*., 2016) and trimmed using the gappy-out method of the trimAl tool (Capella-Gutierrez *et al*, 2009).

### Paralog Filter

There were a number of duplications in the CaSR subfamily. For example, Dipodomys ordii has 116 CaSR sequences. To reduce the number of sequences, human CaSR and other human class C GPCR proteins sequences compiled with CaSR sequences of each taxon and aligned with the MAFFT v7.221 auto algorithm (Yamada *et al*., 2016), and the gappy-out method of the trimAl tool was used to trim the MSA (Capella-Gutierrez *et al*., 2009). The ML tree was built using RAxML-NG v0.9.0 with ML tree search and bootstrapping (Felsenstein Bootstrap and Transfer Bootstrap) parameters (Kozlov *et al*, 2019). Based on the ML tree, proteins that were diverged from the common ancestor of the human CaSR clade were classified as CaSR-likes. Proteins that were clustered with the human CaSR were accepted as CaSRs. After we assigned all proteins to their subfamilies, we built final ML trees for CaSR, GPRC6A, and TAS1Rs. We added human CaSR sequence to the GPRC6A and TAS1Rs subfamilies, and human GPRC6A sequence was added to CaSR subfamily as an outgroup. We aligned subfamily sequences with the MAFFT v7.221 einsi algorithm (Yamada *et al*., 2016) and built the ML trees by using RAxML-NG v0.9.0 with the FTT model parameter (Kozlov *et al*., 2019). We labeled the duplications at each node on the ML trees. Based on the duplications, we manually checked the trees and removed a clade that was a subset of its sister clade by using the ETE toolkit (Huerta-Cepas *et al*., 2016). We took each branch and node length from leaf to root of the tree by using common species in all CaSR, GPRC6A and taste receptor trees to calculate subfamily conservation by using Welch’s t-Test by using the ‘ggstatsplot’ package (Patil, 2021).

### Subfamily Specific Profile HMMs

After we took all receptors from CaSR, CaSR-like, GPRC6A, and taste receptors, we aligned them by using MAFFT v7.221 auto algorithm (Yamada *et al*., 2016). For each subfamily we removed the positions from the MSA that correspond to a gap in the human receptor. Then, we divided the MSA into subfamily alignments. We generated an HMM profile from the gap-removed alignment of each subfamily. The “pnone” option of the HMMER package was used in the HMM profile construction step, and thus the obtained probability parameters are simply calculated by employing observed frequencies. The constructed HMM profiles for any target subfamily, even with the “pnone” option, still hit the sequences from other subfamilies. On the other hand, we aim to obtain profiles such that they can correctly identify the elements of the target subfamily while not hitting any proteins from the remaining ones. To achieve this, we use a position-weighting approach. After determining the weight per position, we update the emission probabilities by taking these weights into account. The weights are computed by a detailed analysis of the conservation dynamics of each subfamily. As the initial step, we defined the target, the closest and the rest groups based on the ML tree. By considering the distance between the root node of each subfamily and the target one, we determined the closest group. The remaining parts of the tree were taken as the rest. To obtain specific HMM profiles for each subfamily, we considered five different scenarios that show variety in terms of the subfamilies in the target, close and rest groups.

- CaSR is the target group, CaSR-likes are the closest group, and GPRC6A and taste receptors (TAS1Rs) are the rest.
- GPRC6A is the target group, TAS1Rs are the closest group, CaSR and CaSR-likes are the rest.
- TAS1R1 is the target group, TAS1R2 is the closest group, and TAS1R3 is the rest.
- TAS1R2 is the target group, TAS1R1 is the closest group, and TAS1R3 is the rest.
- TAS1R3 is the target group, TAS1R1 and TAS1R2 are the closest group and GPRC6A is the rest.

Our main idea is finding the positions that can help discriminate the target family from the others and assigning a high weight to these positions. Similarly, we assign a low weight to the positions that are conserved for all subfamilies since they can cause incorrect hits. The details of how we compute the weight per position is given in Algorithms 1 and 2.

In Step 1 of Algorithm 1, we show how to select the representative amino acids per group and assign a score to each group based on the amino acid frequencies obtained from MSA for any position k. For the target group, the most frequent amino acid, *R*_*T*_, is chosen as the representative one. For the closest and rest groups we first check if the group is composed of a single subfamily or multiple subfamilies. If the group is composed of a single subfamily, we select the most frequent amino acid as the representative and scores are taken as the frequency of this amino acid. Otherwise, we first check whether the most frequent amino acid for at least one element of the group is *R*_*T*_. If it is, *R*_*T*_ is chosen as the representative amino acid for the corresponding group and its frequency is assigned as a group score. If not, the amino acid with the highest frequency for most of the elements is chosen, and the score is computed by taking the average of the frequencies of the representative amino acids in the subfamilies of the corresponding group.

Algorithm 1 – Step 2 shows the details of how we assign type, which represents whether the position can be used to discriminate the proteins in target and compute the initial score for any position k. To calculate the initial score, we defined six different categories and four different position types based on the representative amino acids that we identified in Step 1 of Algorithm 1. Type I, IV, III, II correspond to the positions ordered with respect to the highest to the lowest final weight.

- Category I (lines 12-18): Representative amino acids in the closest and rest groups are gaps. If the representative amino acid in the target group is also gap, then the initial weight type is Type II. Otherwise, the initial weight type is Type I since this position can be used to discriminate between target from close and rest groups.
- Category II (lines 19-25): The representative amino acid in the target group is gap. If the representative amino acid in the closest or rest group is not gap, then the initial weight type is Type I. Otherwise, the initial weight type is Type II.
- Category III (lines 26-56): Representative amino acids in the target, closest and rest groups are different from each other. We check boundary conditions to see whether each group is conserved. If all scores per group are greater than or equal to a predefined threshold value, the position type is I. If any of them is smaller than the threshold, we check whether amino acid substitutions between the representative amino acids are probable with respect to the BLOSUM score. If the BLOSUM score between the representative amino acids of target and close or target and rest is less than the predefined threshold, we again assign Type I. Otherwise, the type is IV.
- Category IV (lines 57-59): The representative amino acids in the target and the representative amino acids in the closest groups are the same. The type is II. These positions are the main reason for the wrong hits, so we assign the lowest weights to the positions in this category.
- Category V (lines 60-70): The representative amino acid in the target group is the same with the representative amino acid in the rest group, but it is different from the representative amino acid in the closest group. If the conservation level of the rest is less than the threshold, the type is III, otherwise, to prevent wrong hits from the rest group to the target, we label the position as Type II.
- Category VI (lines 71-80): The representative amino acid in the closest group is the same with the representative amino acid in the rest group, but the representative amino acid in the target group is different. The type is I if it satisfies the boundary conditions; otherwise, the type is II.

The predefined thresholds *t*ℎ*r*_1_ and *t*ℎ*r*_2_ are chosen by considering the conservation dynamics of target and close&rest, respectively. For example, for CaSR, *t*ℎ*r*_1_ = 0.98 and *t*ℎ*r*_2_ = 0.8. For GPRC6A and TAS1Rs, we took both *t*ℎ*r*_1_ and *t*ℎ*r*_2_ as 0.8. The final threshold that is used to determine whether two amino acids are close to each other in terms of the BLOSUM score, *t*ℎ*r*_*bls*_, is chosen as 0.5. Here, we do not directly use BLOSUM scores of two amino acids, instead we normalize each row of the log odd ratio of the BLOSUM matrix by dividing it by the maximum element of the corresponding row. Thus, the maximum value of the matrix is 1 when the amino acid is conserved and the value decreases with respect to the closeness of substituted amino acids.

After we check each category and assign types to the positions, we calculate the initial scores that will be used to compute the final weight based on the position type. For Types I, III and IV, the initial score is the sum of scores for each group. On the other hand, for Type II, the initial score is computed as one over the sum of individual group scores. Here we aim to assign the lowest weight to Type II since it consists of the positions that can cause wrong hits to target because of similar conservation and amino acid patterns with close and/or rest groups.

After determining the types and initial score of each position, the next and final step is to decide the final weight that will be used to modify the emission probabilities of the default HMM profiles. Details of this process are given in Algorithm 2. The weight of each type will be in the following order from highest to lowest: Type I, Type IV, Type III and Type II. We first order each position in each group from highest to lowest with respect to the initial score. Type I positions take weights greater than equal to *c*_1_ where *c*_1_ = 1 for CaSR, GPRC6A, TAS1R2 and TAS1R3 and *c*_1_ = 1.5 for TAS1R2. The maximum weight of Type IV positions, *c*_2_, is equal to the mean value of the weights of Type I positions (line 4 of Algorithm 2) for all target subfamilies. The maximum weight of Type III positions, *c*_3_, is taken as the minimum weight of Type IV positions for GPRC6A and 0.5 for TAS1Rs. For CaSR, since it is a highly conserved family and Type I and IV positions are much more important compared to the other types, we decreased the maximum weight of Type III positions further by taking the minimum of Type IV over 2. Finally, Type II positions take the lowest score as the highest weight, *c*_4_, is again determined by considering the conservation pattern of the target. We took *c*_4_ = 0.2 for GPRC6A and TAS1R2, *c*_4_ = 0.25 for TAS1R1 and TAS1R3. For CaSR, the highest weight of Type II positions is restricted by the minimum weight of Type III positions over 2. As mentioned earlier, this type includes positions that show similar conservation patterns and cannot be used to distinguish between the elements of the target and other groups.

#### Algorithm 1 Representative amino acid and initial score for position “K”

***Input:*** *Representative amino acid of target subfamily, R*_*T*_*; the frequency of R*_*T*_ *in the target, S*_*T*_*; the most frequent amino acid of subfamily i, (i=1,…,N) and its frequency, a*_*i*_, *F*_*i*_*, respectively; target, close and rest groups, t*, *c*, *and r, respectively; the number of subfamilies in close and rest groups, n*_*c*_ *and n*_*r*_*, respectively; conservation threshold for target and close/rest groups, t*ℎ*r*_1_ *and t*ℎ*r*_2_*; the threshold for Blosum scores, t*ℎ*r*_*bls*._

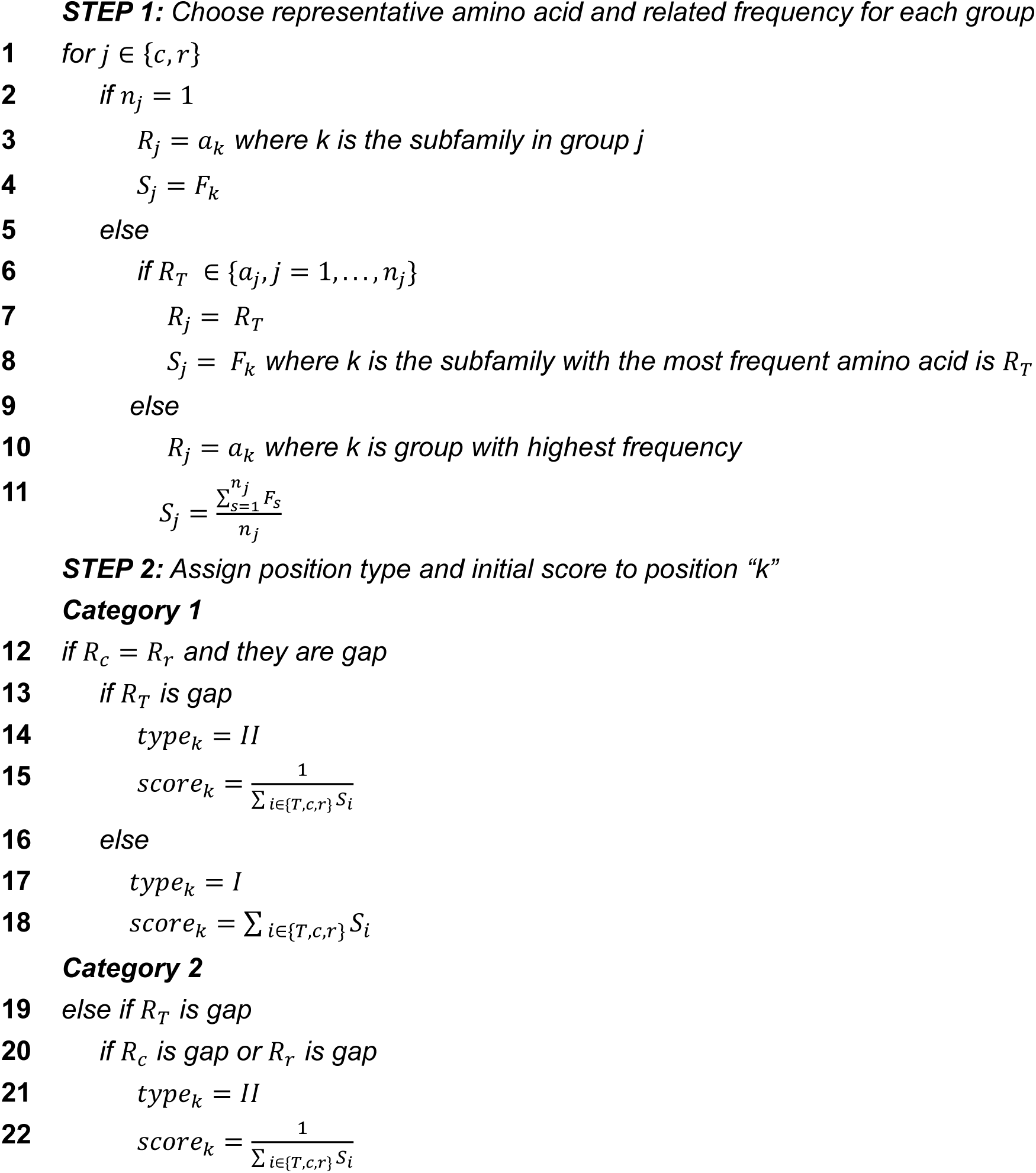
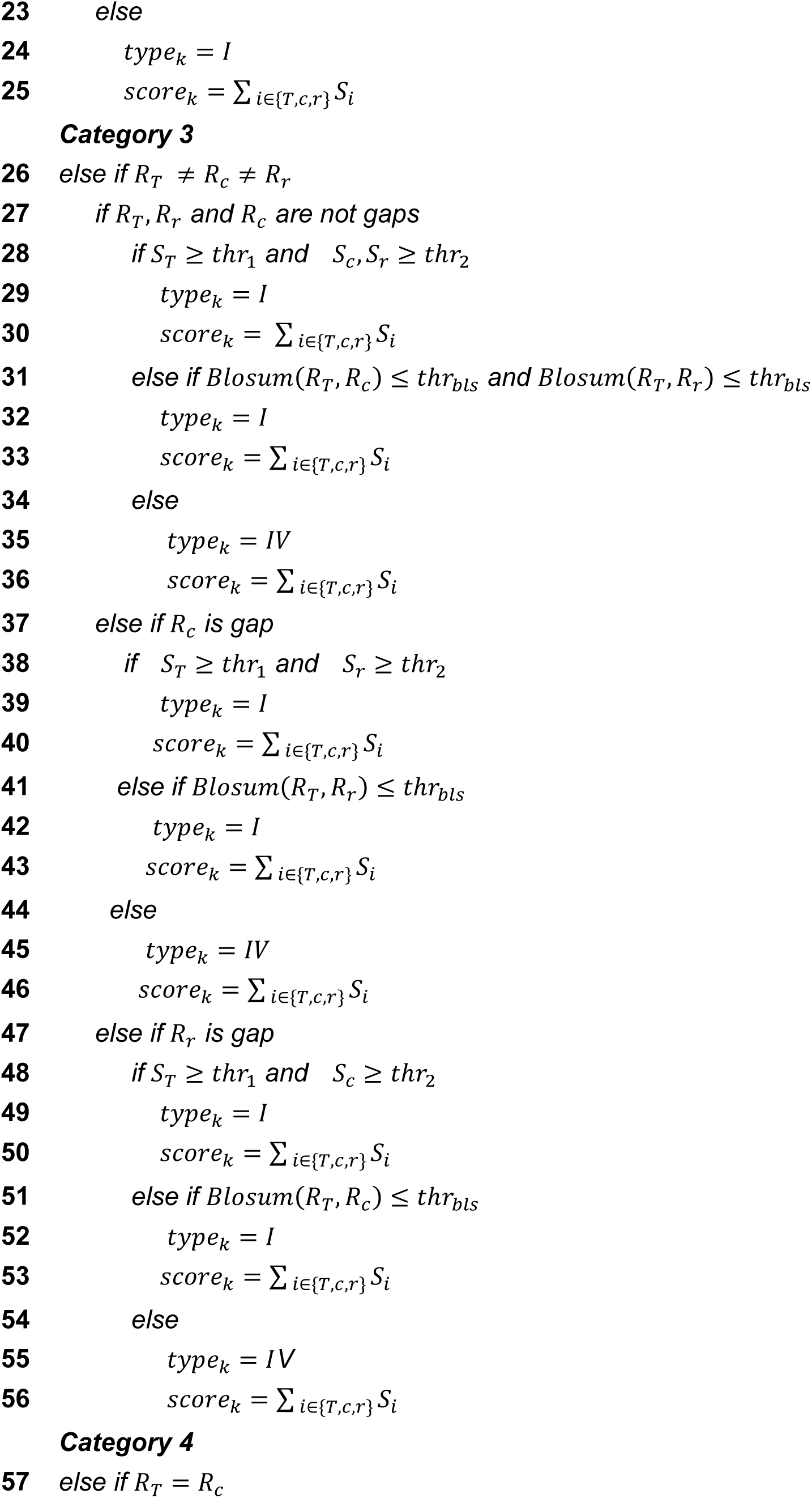
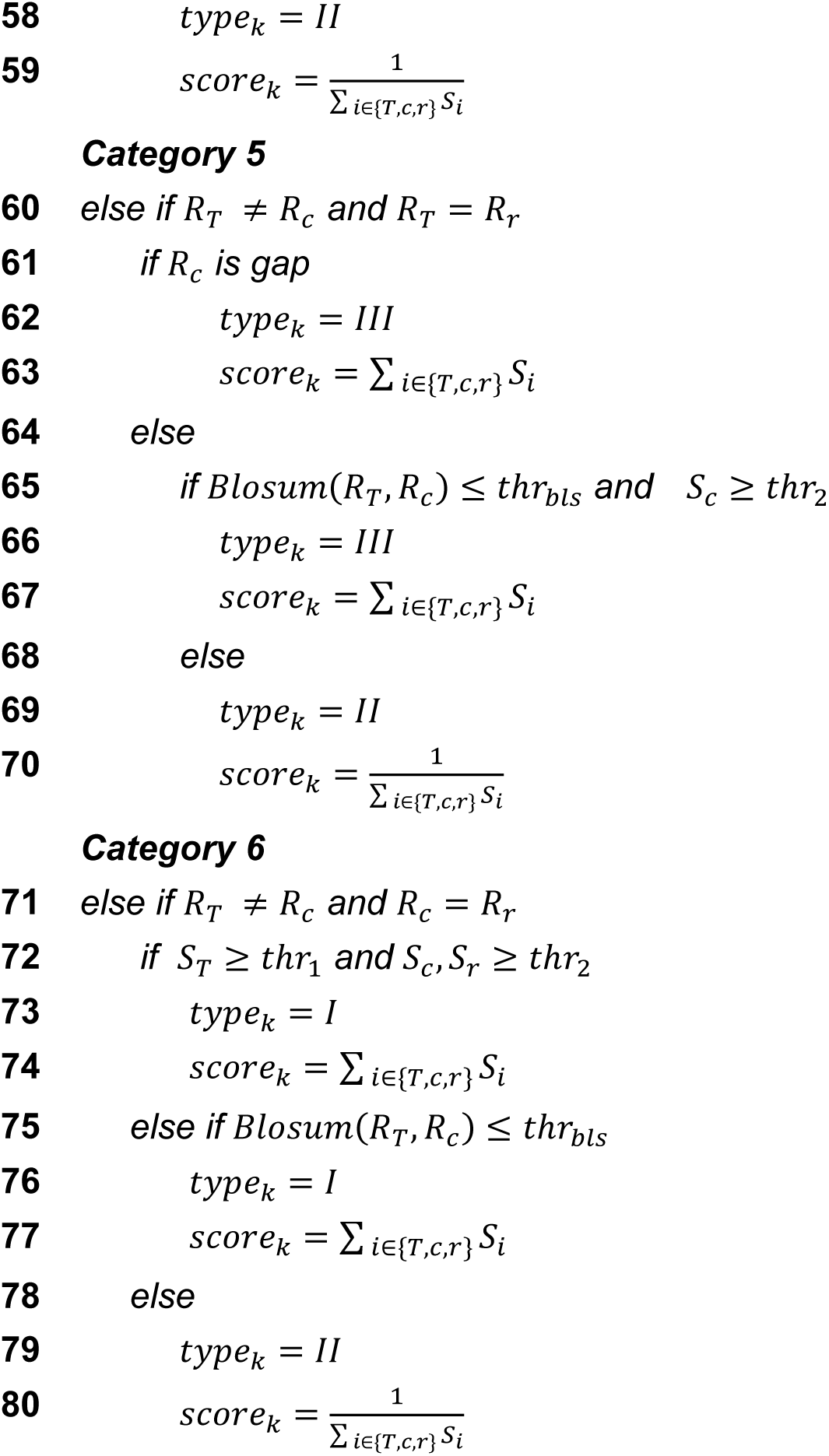

#### Algorithm 2

Compute weight for all positions of target subfamily “S”

***Input:*** *Types for each position k (k=1,…,K), type*_*k*_ *; initial score for each position k of type t, score*_*t*,*k*_*; number of type i positions, n*_*i*_ *where n*_1_ + *n*_2_ + *n*_3_ + *n*_4_ = *K; a predefined constant value as max weight of Type i positions, c*_*i*_*; the target subfamily, S*.

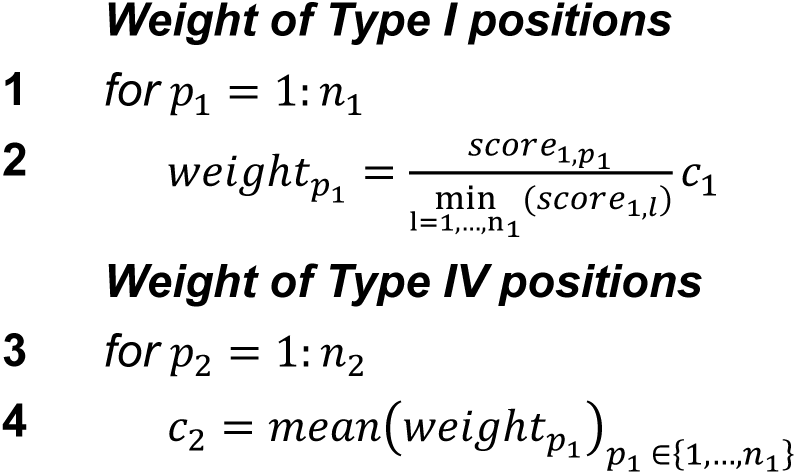
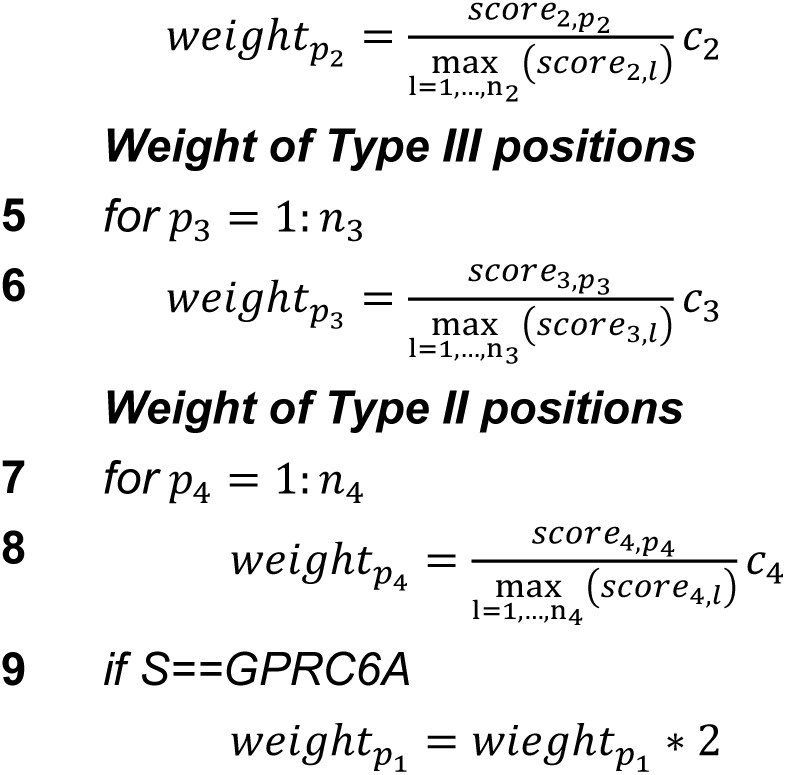

### Subfamily Specific Position Scores

From the alignment we used to make subfamily specific profile HMMs, we randomly selected 264 CaSR like sequences (same number of sequences as CaSRs) and took all CaSR (264 proteins), GPRC6A (242 proteins) and TAS1Rs (TAS1R1 has 210, TAS1R2 has 173 and TAS1R3 has 273 proteins). We built an ML tree by using IQ-TREE multicore version 2.0.6 (Nguyen *et al*, 2015) with automatic model selection (Kalyaanamoorthy *et al*, 2017) (-m MFP) and ultrafast bootstrap (Hoang *et al*, 2018) (-bb 1000) parameters. For CaSR, GPRC6A, and TAS1Rs, we removed the positions from the MSA that correspond to a gap in the human receptor respectively. By using gap removed alignments and the ML tree, we did ancestral sequence reconstructions for each subfamily with IQ-TREE multicore version 2.0.6 with the −m JTT+R10 model parameter (Nguyen *et al*., 2015). We showed specific residues that have a SDP score higher than 5 on the structures. We used the cyro-EM structure of CaSR (PDB:7DTV) and Swiss models (Waterhouse *et al*, 2018) for GPRC6A and taste receptors since they do not have experimental structures. To visualize structures and residues, we used the UCSF Chimera tool (Pettersen *et al*, 2004).

We calculated SDP scores by a method extended from (Bradley & Beltrao, 2019) by considering phylogenetic trees and a phylogeny-based scoring approach, adjPHACT, based on the methodology of the PHACT algorithm. The details of how we compute the SDP score for any position k can be found in Algorithm 3. PHACT computes the tolerance for each amino acid for the query species, which is human, by using a tree traversal approach. By checking the probability differences, PHACT detects the location of amino acid substitutions and compute the weighted sum of positive probability differences based on the distance between the node of change and human. On the other hand, here we aim to determine the acceptability of each amino acid per subfamily. To achieve this, we modify PHACT by starting the tree traversal from the root node and eliminating the node weighting approach. At the end, we have a probability distribution per position for each subfamily, which is computed by considering the independent events. Again, we determine the representative amino acid for the target subfamily by picking the most frequently observed amino acid and its adjPHACT score. For the remaining subfamilies, we keep the adjPHACT score of the representative amino acid of the target and the representative amino acid of the corresponding subfamily. Then, similar to (Bradley & Beltrao, 2019) we check whether the same amino acid is conserved across all subfamilies. On the other hand, our approach differs from (Bradley & Beltrao, 2019) in terms of considering multiple subfamilies and using adjPHACT scores, which employ phylogenetic trees and ancestral reconstruction probabilities. In our approach, we compute the contribution of each subfamily to the SDP score by checking whether the representative amino acid of target has a high adjPHACT score in that subfamily (line 2-7). In the final SDP score for any position k is computed by considering the distance between target and other subfamilies (which is computed by considering the distance between root nodes), the conservation level of the target subfamily in terms of independent amino acid alterations and the individual score coming from each subfamily (line 10).

#### Algorithm 3

SDP Score for position “K”

***Input:*** *Amino acid with the highest adjPHACT score in the target group, a*_*T*_*; the adjPHACT score of a*_*T*_ *in the target,* 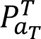 *; adjPHACT score of a_T_ in subfamily I (i=1,…,n),* 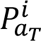*; distance between target subfamily and subfamily I, d_i_; amino acid with the highest adjPHACT score in the subfamily I, a*_*i*_ *; adjPHACT score of a*_*i*_ *in subfamily I,* 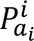.

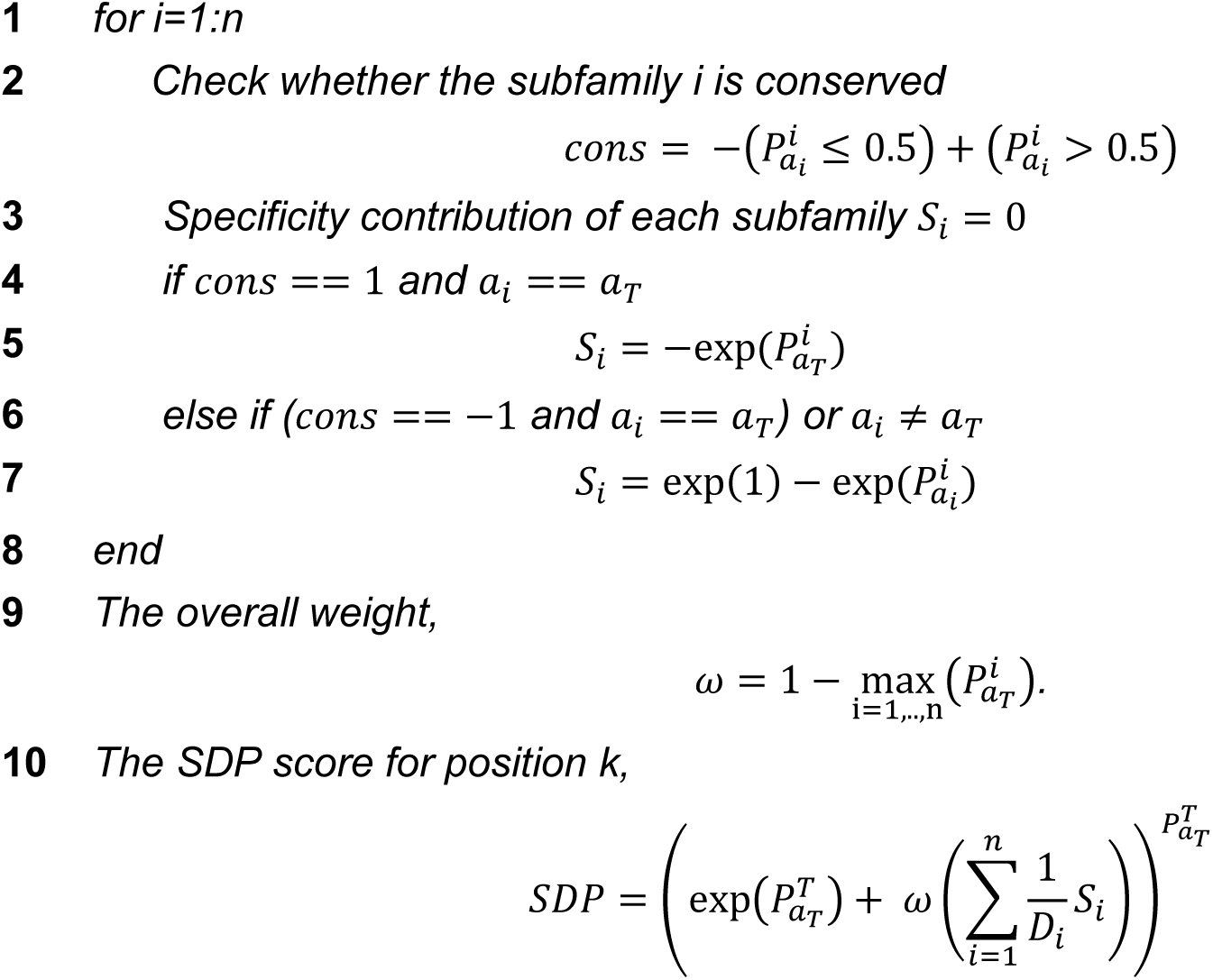

### Evolution of Class C GPCRs

We selected representative sequences from different taxonomic levels for each subfamily and 264 CaSR-like sequences. We aligned them with the MAFFT v7.221 einsi algorithm (Yamada *et al*., 2016). We built the ML tree by RAxML-NG-0.9.0 with the model JTT and transfer bootstrap expectation –bs-metric fbp, tbe parameters (Kozlov *et al*., 2019). We merged the ML trees of CaSR, GPRC6A and taste receptors by checking clades using the ETE toolkit (Huerta-Cepas *et al*., 2016).

### Identification of the CaSR Activation Network

To reveal the network that is important for CaSR activation, we measured changes in contact scores between residues between active and inactive receptor states. For a single protomer, we manipulated the pdb file and the RRCS algorithm (Zhou *et al*., 2019), to be able to detect changes in atom level for identical residues (python codes are provided). We applied a t-test to identify significant changes and chose a p-value threshold of 0.01. Then we later combined atomic-level changes observed within a protomer for each residue to build the activation network. If atoms of a residue are involved in multiple significant changes, this is represented as multiple edges in the network, even if the residue pair is the same.

For the analysis of interactions observed inside the dimerization interface, we used the RRCS algorithm as is and compared residue-residue contact scores between residue pairs upon activation. The important thing to note here is that for every dimer structure we used, we sometimes retrieved two data points due to the symmetrical nature of the dimerization. We again applied the same p-value threshold and identified the important changes observed.

In both of our analyses, we used same set of 7 active state (PDB IDs 7SIL(Park *et al*, 2021b), 7SIM(Park *et al*., 2021b), 7E6T(Chen *et al*., 2021), 7M3G(Gao *et al*., 2021), 7M3F(Gao *et al*., 2021), 7DTT(Ling *et al*., 2021), 7DTV(Ling *et al*., 2021)) and 5 inactive state structures (7SIN(Park *et al*., 2021a), 7E6U(Chen *et al*., 2021), 7M3E(Gao *et al*., 2021), 7M3J(Gao *et al*., 2021), 7DTW(Ling *et al*., 2021)) human CaSR structures.

## Machine Learning

### Dataset and Feature Preparation

To predict the consequence of a substitution in human CaSR, we used a gradient boosting-based machine learning algorithm, XGBoost (Chen & Guestrin, 2016). We used the XGBoost library for R (Chen *et al*, 2015) to train our model. We selected a total of 337 LoF and GoF mutations from the literature (Gorvin, 2019) to train our model. Since we used conservation scores as features to train our model, we divided subfamily alignments and mutations randomly as 80% training and the remaining 20% test data before creating feature matrices to prevent information leakage. 25% of the training data was randomly picked as the validation data five times for cross-validation. For each dataset split we used the sklearn train test split model with stratify option to keep the LoF to GoF ratio almost the same in the datasets (Pedregosa *et al*, 2011). We calculated the conservation score of the reference amino acid and the substituted amino acid in human CaSR in each subfamily. The reference and the substituted amino acids were represented by BLOSUM62-encoded matrices. Amino acid physico-chemical feature values Zimmerman polarity (Zimmerman *et al*, 1968), average flexibility (Bhaskaran & Ponnuswamy, 1988), Dayhoff (Barker *et al*, 1978), average buried area (Rose *et al*, 1985), Doolittle hydropathicity (Kyte & Doolittle, 1982), atomic weight ratio (Grantham, 1974), molecular weight, and bulkiness (Zimmerman *et al*., 1968) from the ProtScale database (Walker, 2005); and domain information of the reference amino acid were used as other features. We normalized the physico-chemical feature values prior to model training. We repeated the whole random dataset splitting and feature preparation procedure 50 times to obtain more robust results.

### Model Selection and Parameter Tuning

We picked the model parameters for each replication by applying a 5-fold cross-validation technique to the training set. We tuned the model parameters step-by-step using the same validation sets for each parameter to decrease the time complexity. We used the following order of model parameters, so that the parameter that has the highest impact on model outcome was tuned first: Eta and nrounds, gamma, maxdepth, subsample, colsample bytree, min child weight, lambda, alpha. We selected the maxdepth as 2, the minimum maxdepth value to prevent overfitting. We chose eta, gamma, colsample bytree, subsample, min child weight from the sets 0.00001,0.00002,…, 0.001,0,0.1,0.2,…,0.5, 0.5,0.55,…,1, 0.5,0.55,…,1, 1,2,…,6 respectively. We selected regularization parameters lambda and alpha from the set 0, 1e-4, 1e-3, 1e-2, 1e-1, 1, 10, 100. We set the nrounds parameter to 200.

### Performance Metrics

We used the area under the receiver operating characteristic curve (AUROC) and the area under the precision-recall curve (AUPR) to evaluate the performance of our prediction model. AUROC and AUPR are performance measures that are widely used to evaluate the performance of binary classification problems. The higher the AUROC and AUPR, the better the model distinguishes classes. To understand how our model makes predictions, we used SHAP (SHapley Additive exPlanations) values. Shap values give an estimate of how much ach feature contributed to the prediction of the model made (Lundberg & Lee, 2017). We calculated SHAP values for our final model trained by all samples by the using R shapviz package (Mayer, 2023).

### Predictive Performance

After we evaluated the performance of our machine learning algorithm over 50 replications, we used the whole dataset to train the model that we used to make predictions for every possible mutation in human CaSR. We selected model parameters by using the 5-fold cross-validation technique on the whole dataset. To create a new test dataset, we took subfamily alignments of the species from the new Uniprot dataset that did not exist in the training data. We eliminated amino acids that are observed in the CaSR alignment as neutral. In each position, we predicted the GoF or LoF class for any substitution. We did a literature search to find new clinical cases that cause either GoF or LoF mutations. We reported our predictions in the table.

## Data Availability

The open-source code is available at our GitHub repository: https://github.com/CompGenomeLab/CaSR

## Acknowledgments

This study was supported by EMBO Installation Grant no:4163 funded by TÜBİTAK (to OA). Ogun Adebali is supported by the BAGEP program of the Science Academy - Türkiye, and the TÜBA-GEBİP program of the Turkish Academy of Sciences.

## Notes

### Competing Interest Statement

The authors have declared no competing interest.

### Summary of Updates

Updated Version

